# Mucosal BCG vaccination reprograms lung interstitial macrophages and enhances antimicrobial defense in mice

**DOI:** 10.64898/2026.02.03.703470

**Authors:** Aaron James Forde, Mara Esposito, Emanuela Kerschbamer, David Schreiner, Eduardo Moreo, Nikos Pantouloufos, Tiphaine MN Camarasa, Jean de Lima, Habiba Soliman, Maike Erber, Claire E. Depew, Wadschma Naderi, Carolyn G. King

## Abstract

Bacille Calmette–Guérin (BCG) is the only licensed vaccine against tuberculosis (TB) but provides inconsistent protection against disease. Studies in non-human primates show that intravenous BCG confers superior protection compared to intradermal vaccination, yet how different vaccination routes reprogram lung immunity in mice - the primary model for mechanistic studies - remains incompletely defined at single cell or spatial resolution. Moreover, while lung macrophages are key effectors during *Mycobacterium tuberculosis* infection, most studies have focused on alveolar macrophages, leaving interstitial macrophage responses to vaccination largely unexplored. Here, we define the cellular and spatial immune landscape shaped by three BCG vaccination routes in mice using flow cytometry, single-cell RNA sequencing, and spatial transcriptomics. Comparing intratracheal (IT), intravenous (IV), and subcutaneous (SC) vaccination, we demonstrate that mucosal IT delivery uniquely reprograms interstitial macrophages, establishes spatially organized immune hubs with CD4 T cells, and provides superior protection against infectious challenge. These findings identify IM as key mediators of mucosal vaccine-induced protection and provide a framework for the rational optimization of vaccines against respiratory pathogens.

## Introduction

Tuberculosis (TB) remains one of the world’s deadliest infectious diseases, causing an estimated 10.8 million cases and 1.25 million deaths in 2023, with the highest burden in resource limited low- and middle-income countries (1). Despite widespread use for over a century, Bacillus Calmette-Guérin (BCG) remains the only licensed vaccine against TB and provides inconsistent protection against pulmonary disease. A deeper understanding of the immune mechanisms underlying durable protection is urgently needed.

Lung macrophages are among the first immune cells to encounter *Mycobacterium tuberculosis* (Mtb), the causative agent of TB, following inhalation. Two major macrophage subsets inhabit the lung: alveolar macrophages (AM), which reside in the airspaces, and interstitial macrophages (IM), which occupy the lung parenchyma (2). These macrophages differ in origin, metabolic state and antimicrobial capacity. AM are the earliest responders to Mtb, and accumulating evidence suggests that prior vaccination can metabolically and epigenetically reprogram them to restrict Mtb growth, ultimately reducing early bacterial burden and shaping the trajectory of disease progression (3,4). In comparison, the precise nature of IM rewiring following BCG vaccination is less well understood. Surface marker expression levels of CD206, CD11c, MHCII, Lyve1, CX3CR1 and CD169 have been used by different groups in an effort to further describe IM heterogeneity in the naive lung (5–7). Additional work to better characterize different subgroups of IM has shown a distinct functional separation based on expressed chemokines (8). However, how IM might be modulated during infection remains largely unknown. Importantly, IM are not a static population: upon niche depletion, Ly6c high monocytes undergo local differentiation into IM in an M-CSF, MafB and TGFβ dependent manner, suggesting that the IM compartment retains plasticity and could be exploited by vaccination (9,10). Moreover, evidence suggests that the role of IM during pulmonary infection may be context dependent. For example, IM have been shown to be permissive to *Klebsiella pneumoniae* and depletion of IM in a fungal infection model resulted in significantly reduced pathogen load (11,12). In contrast, mouse models of Mtb infection have shown that IM are more restrictive to bacterial growth as compared to AM (13,14).

Although BCG is typically administered intradermally, recent studies demonstrate that vaccination route significantly impacts protective immunity against Mtb. In non-human primates, intravenous BCG vaccination conferred robust protection following Mtb challenge, induced markedly higher frequencies of antigen-responsive CD4 and CD8 T cells in blood, lung parenchyma, and bronchoalveolar lavage fluid compared to intradermal vaccination, and conferred robust protection following Mtb challenge (15,16). Subsequent studies showed that high-dose IV BCG drives an influx of polyfunctional T cells and recruited macrophages into airways, with both CD4+ and CD8α+ T cells required for protection (15,17). Similar route dependent effects have been observed in mice. Intranasal but not subcutaneous BCG vaccination activates AM, enhancing metabolic and activation gene signatures and improving bacterial control after both Mtb and *Streptococcus pneumoniae* challenge (18). More recently, IV BCG was shown to generate a CX3CR1-high CD4+ effector memory T cell subset capable of mediating cross protection against influenza via IFNγ dependent activation of AM (19).

Collectively, these findings suggest that vaccination route shapes immune responses in the lung, with protection strongly correlating with enhanced T cell and macrophage activation. However, studies to date have focused primarily on AM while the impact on IM biology remains largely unexplored. Notably, IM express high levels of the anti-microbial effector NOS2 during Mtb infection in mice, suggesting a potential role in early bacterial control (14). Whether vaccination, particularly via alternative routes, enhances IM antimicrobial capacity remains unknown. This gap is especially relevant given growing interest in mucosal vaccination strategies which may directly engage pulmonary macrophages at the site of pathogen entry and negate safety concerns associated with IV BCG injection.

To directly compare how vaccine delivery route shapes lung macrophage biology, we immunized mice with BCG via SC, IV, or mucosal IT administration. High-dimensional flow cytometry and single-cell RNA sequencing (scRNA-Seq) revealed that IT vaccination selectively expands a distinct IM subset enriched for metabolic, antimicrobial, and type I/II IFN-response programs that were not observed after SC or IV delivery. Consistent with these findings, IT BCG elicited the strongest protection in Mtb rechallenge experiments. Integration of cell-cell interaction analyses and spatial transcriptomics uncovered potential IM - CD4 T cell interactions and demonstrated that activated IM localize to distinct regions of the lung following IT BCG. Together, these findings identify IM reprogramming as a previously underappreciated component of mucosal BCG immunity and highlight mucosal vaccination as a promising strategy to harness lung macrophages for improved protection against Mtb.

## Results

### IT BCG supports enhanced IM activation

Given that lung macrophages are among the first immune cells to encounter pulmonary Mtb, we focused on how the route of BCG vaccination differentially rewired these cells. Wild type (WT) mice were vaccinated with SC, IT, or IV BCG, and after 8 weeks lungs were harvested and analyzed by high dimensional flow cytometry using a gating strategy designed to identify AM, IM as well as other myeloid cells (Supplementary Fig. 1 A) (6).

The frequency of AM was reduced after IT and IV vaccination compared to naïve mice but the absolute numbers remained unchanged (Fig. 1 A). Analysis of typical macrophage activation markers including CD80, CD86 and MHCII revealed increased expression after IT and IV, but not SC, vaccination in AM (Fig. 1 B). CD38 has recently been described as a marker of macrophages which possess an enhanced ability to control Mtb (20). Following IT and IV vaccination the frequency of CD38 positive AM was increased (Fig. 1 C). In contrast, SC BCG did not lead to enhanced CD38 expression on AM (Fig. 1 C). The frequency of NOS2 positive AM increased by a small but significant extent following IT and IV, but not SC, vaccination (Fig. 1 D). We next examined the AM phenotype at an earlier time point of 4 weeks post vaccination. Following IT and IV vaccination the expression of CD80, CD86 and MHCII was increased on AM, as was the frequency of CD38 positive cells. Similar to the 8 week time point, SC BCG did not enhance the expression of the analysed markers (Supplementary Fig. 1 B-D). At a later timepoint of 12 weeks post vaccination the increased expression of CD86 and MHCII was maintained in IT and IV vaccinated groups as compared to naive or SC vaccinated mice (Supplementary Fig. 1 E). Increased frequencies of CD38 or NOS2 positive AM were only observed in the IT vaccinated group (Supplementary Fig. 1 F, G).

**Figure 1:**
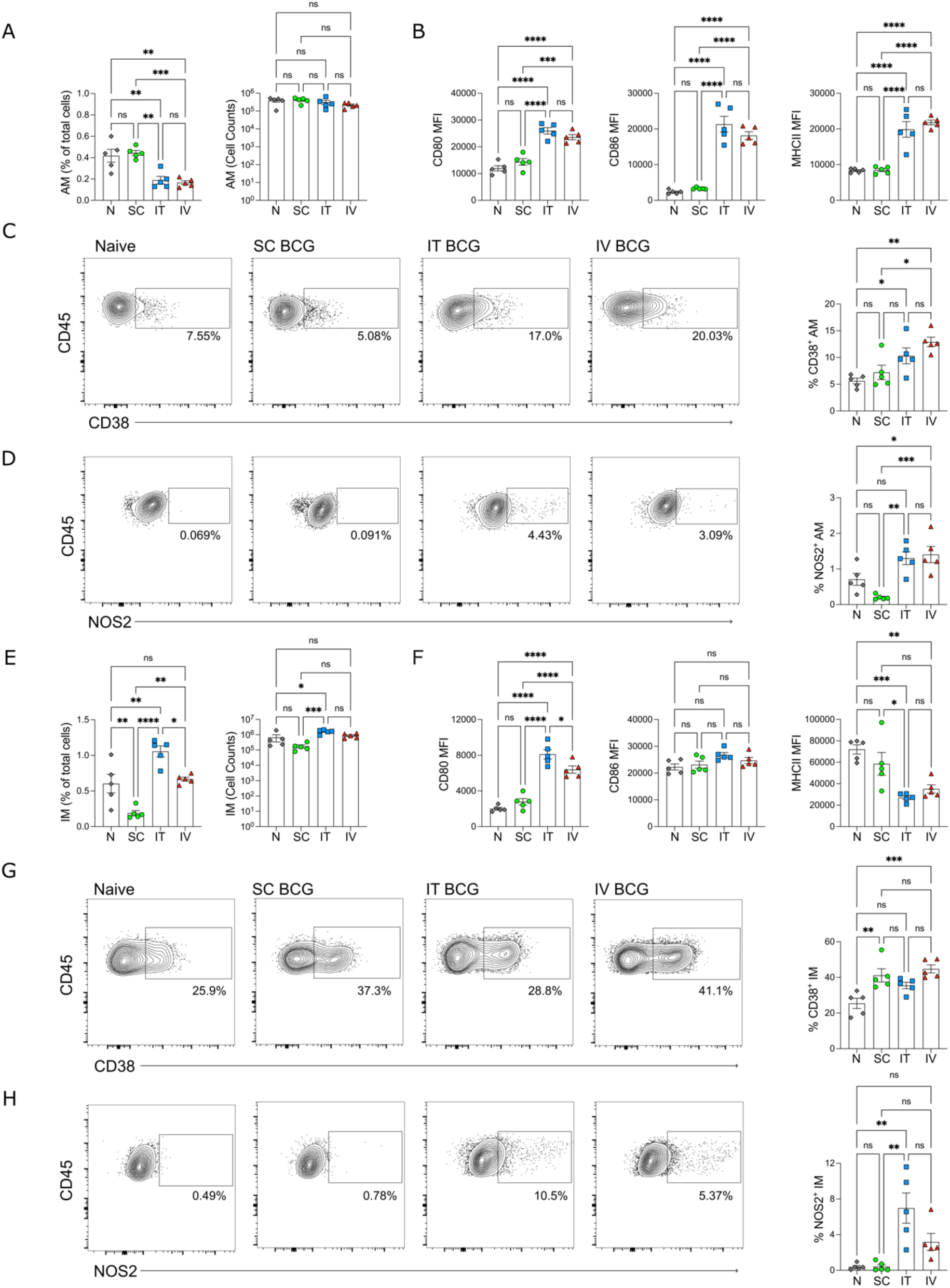
IT BCG supports enhanced IM activation. **(A)**: Frequency of total lung cells and absolute numbers of AM 8 weeks after SC, IT and IV BCG vaccination. **(B)**: Mean Fluorescence Intensity (MFI) of CD80, CD86 and MHCII expressed by AM after BCG vaccination. **(C, D)**: Representative plots and quantification of CD38 or NOS2 positive AM 8 weeks after BCG vaccination. **(E)**: Frequency of total lung cells and absolute numbers of IM 8 weeks after SC, IT and IV BCG vaccination. **(F)**: MFI of CD80, CD86 and MHCII on IM 8 weeks vaccination. **(G, H)**: Representative plots and quantification of CD38 or NOS2 positive IM 8 weeks after BCG vaccination. N: Each datapoint indicates one individual mouse. One-Way ANOVA with Tukey’s multiple comparison test was used to determine statistical significance.

Within the IM compartment we observed an increase in both cell frequency and absolute numbers 8 weeks after IT BCG (Fig. 1 E). Analysis of activation markers revealed increased expression of CD80, no change in CD86 and downregulation of MHCII after IT and IV BCG (Fig. 1 F). In contrast to AM, the frequency of CD38 positive IM increased after all routes of vaccination (Fig. 1 G), while NOS2 expression was significantly increased only after IT BCG (Fig. 1 H), suggesting route-specific enhancement of antimicrobial effector capacity. At 4 weeks post vaccination CD80 expression was increased after IT and IV BCG, however CD86 and MHCII levels were decreased (Supplementary Fig. 1 H). Furthermore, IT and IV BCG also increased the frequencies of both CD38 and NOS2 positive cells (Supplementary Fig. 1 I, J). By 12 weeks post vaccination IM from IT BCG mice maintained elevated CD80 expression, whereas the expression of the other analyzed markers appeared to be returning to baseline (Supplementary Fig. 1 K-M). Together, these data indicate that IT and IV BCG exert significant alterations to the phenotype of both AM and IM, while SC BCG has a minimal impact.

Since viable BCG is present up to 12 weeks post IT and IV vaccination (Supplementary Fig. 1 N), we next wondered if the continued presence of bacteria was required for driving the expression of CD80, CD86 and MHCII by AM or IM. To examine this, cohorts of WT mice were vaccinated using SC, IT or IV routes for a period of 8 weeks. Next, the animals were administered a cocktail of antibiotics (rifampicin and isoniazid) in the drinking water for 4 weeks. Following a rest period of 2 weeks, the lungs were harvested and analysed using flow cytometry. We observed similar expression of CD80, CD86 and MHCII by AM after all routes of vaccination in antibiotic treated and untreated mice (Supplementary Fig. 1 O). In the case of IM, antibiotic treatment did not influence the expression of these markers, with the exception of increased MHCII expression in IT-vaccinated mice (Supplementary Fig. 1 P).

We next used scRNA-Seq to more comprehensively characterize the transcriptional programs underlying changes in macrophage phenotype. CD45 positive cells were sorted from the lungs of naive mice as well as those that had been vaccinated with SC, IT or IV BCG 8 weeks prior. The cell suspensions were stained with CITE-seq antibodies to facilitate the identification of the various immune cell populations. We began our analysis by focusing on myeloid cells and using surface markers and known marker genes, we identified AM, IM, DCs and monocytes (Fig. 2 A, Supplementary Fig. 2 A-D). It was notable that among these populations SC BCG appeared to have a minimal impact on the transcriptome of AM and IM (Supplementary Fig. 2 E). In contrast IT and IV BCG resulted in alterations to the transcriptomic profile of both cell types (Supplementary Fig. 2 F, G).

**Figure 2:**
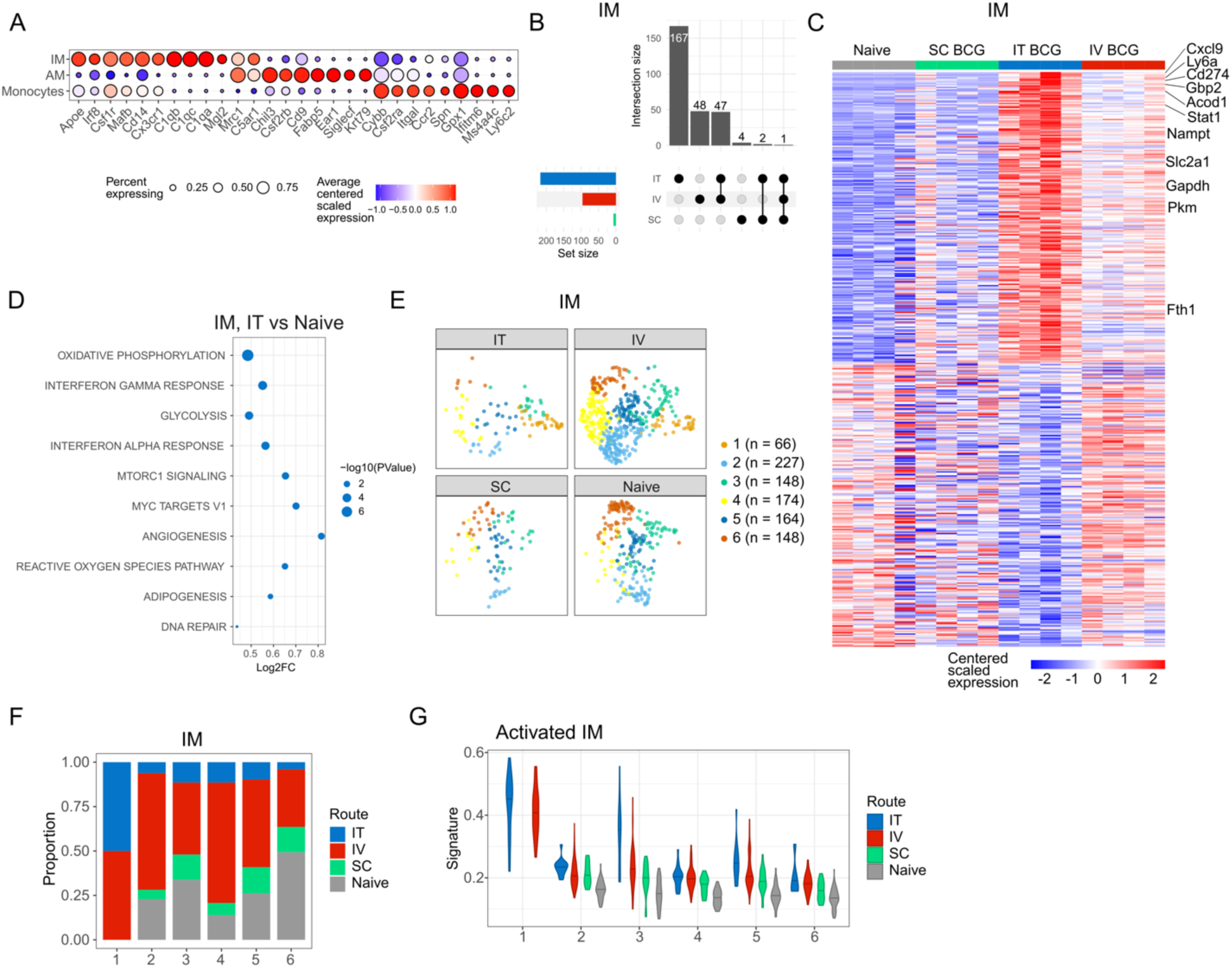
IT BCG transcriptionally rewires IM. **(A):** Expression of signature genes used to identify AM, IM and monocytes in the single cell RNA-Seq dataset. **(B):** Upset plot depicting the number of genes upregulated by IM after each route of BCG vaccination as compared to naïve cells. **(C)**: Heatmap depicting differentially expressed genes by IM following IT BCG vaccination. **(D)**: Pathway enrichment analysis performed on the genes upregulated by IM 8 weeks after IT BCG vaccination. **(E):** UMAP displaying IM clusters that were present in naive mice and after SC, IT or IV BCG vaccination. **(F):** Proportion of IM clusters described in (E). **(G)**: Violin plot depicting the activated IM expression signature score in each IM cluster.

Pseudobulk differential expression analysis further revealed that IV BCG led to the most upregulated genes in AM as compared to naïve controls (Supplementary Fig. 2 H), while IT BCG-treated AM showed increased expression of antigen presentation genes *(H2-K1*, *H2-Ab1*) as well as the chaperone protein *Cd74* (Supplementary Fig. 2 J). We next focused our analysis on IM after vaccination and observed that IT BCG had the most profound effect in terms of the number of upregulated genes (Fig. 2 B). Within these upregulated genes were a number of interferon targets such as *Cxcl9*, *Gbp2, Stat1* and *Cd274* (Fig. 2 C). In addition, IM from IT BCG vaccinated mice expressed several metabolic genes including *Gapdh*, *Nampt*, *Pkm*, *Fth1, Slc2a1* as well as *Acod1* (Fig. 2 C). In line with this, gene set enrichment analysis revealed increased activity in pathways associated with IFNγ response, OXPHOS and glycolysis in IM from IT vaccinated mice (Fig. 2 D). Given that the aforementioned analysis was performed on all IM, we wondered if only a subset of IM were responsible for driving the expression of these pathways. We identified 6 clusters of IM in naïve and vaccinated mice of which cluster 1 was present only after IT and IV vaccination (Fig. 2 E, F). In order to examine if the IM within cluster 1 were the cells primarily activated following BCG vaccination, we created a gene expression signature score (named activated IM) based on the genes upregulated after IT BCG. This signature score was most highly expressed by the IM present in cluster 1 (Fig. 2 G) indicating that this subset was primarily activated after IT BCG.

### CD4 T cells likely produce IFN**γ** after all routes of BCG vaccination

Given that IM from IT vaccinated mice showed enrichment for IFNγ-response genes (Fig. 2 C-D), we next sought to identify which immune cells were producing IFNγ. To do this we examined the non-myeloid immune cells within our single cell data set and based on marker genes and surface protein expression identified B cells, NK cells, as well as CD4 and CD8 T cells (Supplementary Fig. 3 A, B). Within these lymphocyte populations *Ifng* expression was highest in the CD4 T cell compartment (Supplementary Fig. 3 C). We therefore focused on examining route dependent differences in CD4 T cells following BCG vaccination. 5 clusters of CD4 T cells were identified in naïve and BCG vaccinated mice (Supplementary Fig. 3 D). Cluster 5 was present predominantly in unvaccinated mice and were likely unactivated cells. Clusters 2 and 4 expressed genes such as *Sell*, *Ccr7* and *S1pr1,* these are expressed by naïve and central memory CD4 T cells (Supplementary Fig. 3 E, F). Despite sharing the expression of the above-mentioned genes, clusters 2 and 4 differed in their distribution among the experimental groups. Specifically, cluster 2 was present only after IT and IV BCG while cluster 4 was present in SC vaccinated mice (Supplementary Fig. 3 G). Genes higher in cluster 2 than cluster 4 suggest more active translational machinery: *Eif3f* (a component of the translation initiation complex), the transcription factor *Klf13*, *Chd3* (involved in chromatin remodeling), *Tut4* (involved in mRNA decay) and *Pnrc1* (Supplementary Fig. 3 E, F).

Cluster 1 was present in all conditions but at a higher proportion in IT and IV vaccinated mice (Supplementary Fig. 3 G). This cluster resembled migrating CD4 T cells (*Maf* and *Cxcr3*) and a small proportion of Tfh cells (*Bcl6*, *Pdcd1*, *Izumo1r*, *Cxcr5*) (21). We next performed pseudobulk differential expression analysis and compared how the CD4 T cells within cluster 1 differed after each route of vaccination. After IT and IV vaccination, cells in cluster 1 had a higher expression of both *Socs1* and *Jund* as compared to SC BCG, potentially indicating a more regulatory phenotype (Fig. 3 A, B). In addition, *Tgfb* expression was increased after IT and IV as compared to SC BCG. Beyond its role in tissue remodeling, TGFβ signaling has recently been described to mediate the transition of monocytes to IM in the lung (10) and is required for AM survival (2,22). Interestingly, when comparing cells within cluster 1 in the IT and IV groups, we observed increased expression of *Hk2* in the IT group which may be indicative of enhanced metabolic activity. These cells also exhibited higher expression of *Cxcr4* (a Tfh marker) and *Cxcr6* (a Th1 marker), genes which are involved in the homing to, and retention in, inflamed tissue (Fig. 3 C) (23,24) .

**Figure 3:**
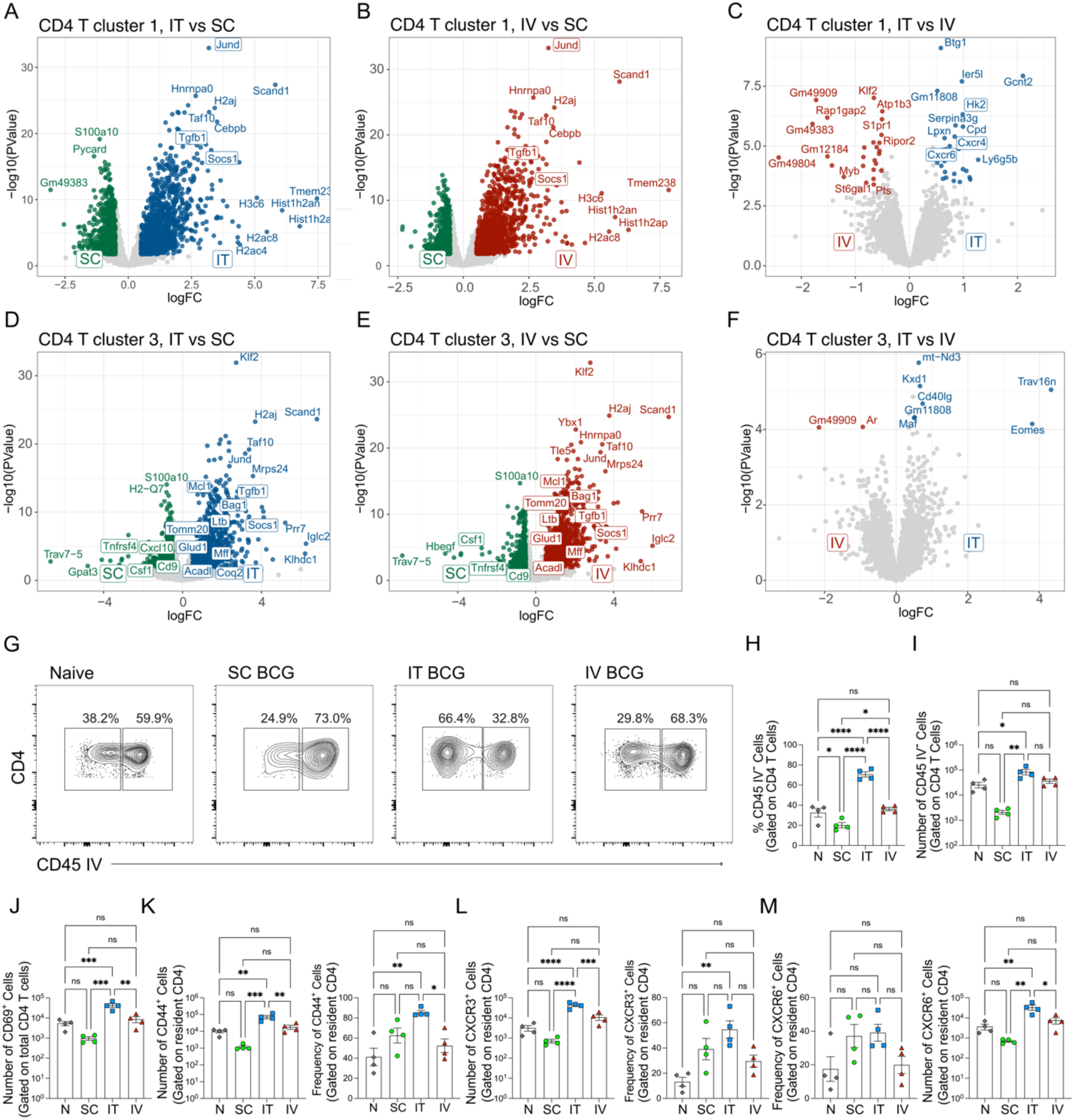
CD4 T cells likely produce IFNγ after all routes of BCG vaccination. **(A-C)**: Volcano plots depicting DEGs in cluster 1 CD4 T cells IT vs SC (A), IV vs SC (B) or IT vs IV (C). **(D-F)**: Volcano plots depicting DEGs in cluster 3 CD4 T cells IT vs SC (D), IV vs SC (E) or IT vs IV (F). **(G)**: Representative plots depicting the proportions of CD45 IV positive and negative CD4 T cells in naive mice or in animals vaccinated with SC IT or IV BCG 8 weeks prior. **(H, I)**: Frequency (H) and absolute number (I) of CD45 negative CD4 T cells in naive and vaccinated mice. **(J)**: Number of CD69 positive resident CD4 T cells in naive mice and those vaccinated 8 weeks prior. **(K-M)**: Frequency and absolute numbers of CD45 negative CD4 T cells expressing CD44 (K), CXCR3 (L) and CXCR6 (M). For Fig. 3 H-M, N: Each datapoint indicates one individual mouse. One-Way ANOVA with Tukey’s multiple comparison test was used to determine statistical significance. For volcano plots the coloured dots represent significantly differentially expressed genes with FDR < 0.05 and |logFC| > 0.5 (dark grey for Naive, green for BCG SC, blue for BCG IT, red for BCG IV). Light grey dots represent non-significant genes.

CD4 T cells within cluster 3 expressed genes such as *Cx3cr1*, *Zeb2*, *Il18rap* and *Ifng* (Supplementary Fig. 3 E, F). This cluster resembled a recently described effector memory subset of CX3CR1^hi^ CD4 T cells that are induced following IV BCG (19). To confirm this apparent similarity, we generated a gene expression signature score (named activated CD4 cells) based on the aforementioned publication and overlaid it onto our single cell dataset. As expected, cluster 3 had the highest signature expression score followed by cluster 1 (Supplementary Fig. 3 H, I). Notably, cluster 3 CD4 T cells were present at similar proportions across all vaccination routes (Supplementary Fig. 3 G), prompting us to investigate whether functional differences might be more apparent at the transcriptional level. When comparing IT and SC routes we observed that the cells from the IT mice expressed metabolic genes such as *Glud1*, *Acadl*, *Coq2*, *Mff*, and *Tomm20* suggesting an enhanced oxidative metabolism typical of memory cells. Moreover, they expressed genes indicative of a regulatory profile like *Tgfb1, Ltb* and *Socs1* as well as the anti-apoptotic genes *Mcl1* and *Bag1* (Fig. 3. D). In contrast, CD4 T cells within cluster 3 from SC BCG mice appeared to be in a more inflammatory state. These cells expressed *Cd9* (a tetraspanin associated with immune synapse formation), *Cxcl10* (involved in T cell recruitment), *Csf1* (indicating ongoing communication with monocytes or macrophages) and *Tnfrsf4* (OX40, a costimulatory receptor). Similar gene expression patterns were observed when cluster 3 CD4 T cells were examined in the IV versus SC groups (Fig. 3 E).

To complement these data we performed additional flow cytometry experiments with a panel targeting CD4 T lymphocytes (Supplementary Fig. 3 J). 8 weeks after vaccination the absolute cell number of total CD4 T cells was increased in IT BCG mice (Supplementary Fig. 3 K, L). In addition, the frequency and number of tissue resident CD45 IV negative CD4 T cells was increased after IT BCG (Fig. 3 G-I). In line with this we observed an increase in the number of total CD4 T cells expressing the residency marker CD69 (Fig. 3 J). Further, CD45 IV negative CD4 T cells from IT BCG mice increased expression of CD44, CXCR3 and CXCR6, collectively indicating enhanced activation and potential tissue retention (Fig. 3 K-M).

### IM-CD4 T cell crosstalk following BCG Vaccination

Having identified route specific differences in both IM and CD4 T cell phenotypes, we next performed NicheNet analysis to understand if these cell types were directly interacting. For this analysis we separated CD4 T cells from cluster 3 (labeled activated CD4 T cells) from the other clusters (labeled as CD4 T cells). This confirmed that activated CD4 T cells were the primary potential source of IFNγ for IM after all vaccination routes (Fig. 4 A-C) and that this was signaling via *Ifngr1* and *Ifngr2* on IM (Supplementary Fig. 4 A-C). Next, we wondered if IM were reciprocally signaling to CD4 T cells. NicheNet analysis predicted that after IT BCG, IM signal to CD4 T cells primarily via the chemokines *Ccl3*, *Cxcl10* and *Cxcl16* (Fig. 4 E). *Cxcl16* was predicted to signal via *Cxcr6* (Supplementary Fig. 4 E). The predicted *Cxcl16*/*Cxcr6* signaling axis was consistent with our flow cytometry data showing increased CXCR6 expression on CD4 T cells after IT vaccination (Fig. 3 N-P). IM from IV BCG mice were similarly providing *Ccl3* and *Cxcl16* (Fig. 4 F). In contrast to IT and IV vaccination, CD4 T cells from SC BCG mice were not receiving these ligands from IM and instead were receiving a distinct set of signals including *Il-1b,* as well as *Ifng* from activated CD4 T cells (Fig. 4 D). Collectively these data suggest that mucosal IT vaccination establishes a bidirectional IM-CD4 T cell circuit such that IM derived CXCL16 promotes the recruitment of CD4 T cells to the lung, which in turn provide IFNγ to sustain IM activation.

**Figure 4:**
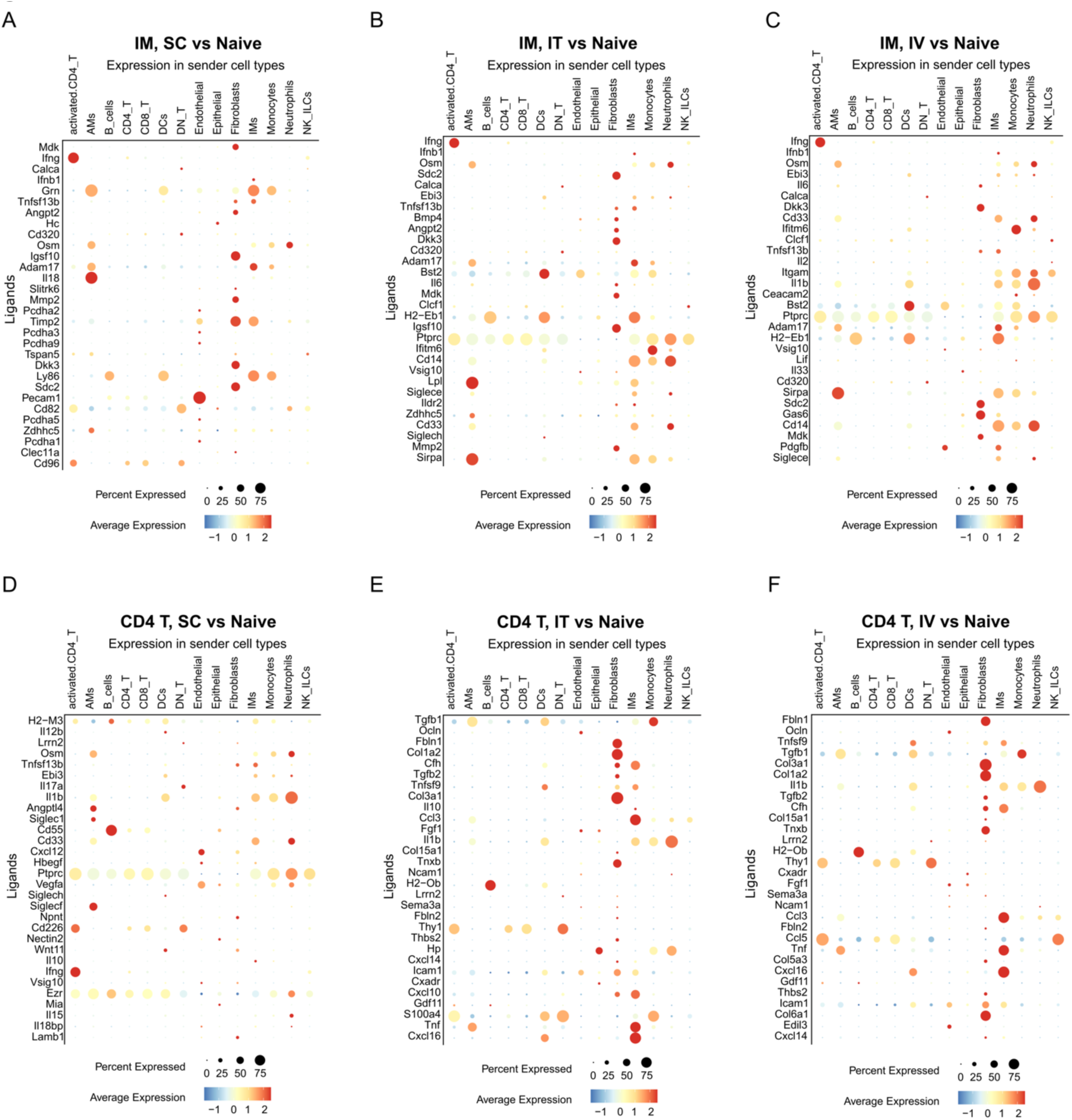
IM-CD4 T cell crosstalk following BCG Vaccination. **(A-C)**: NicheNet analysis showing the expression of ligands received by IM (compared to Naive) in the sender cell type after each route of vaccination. **(D-F)**: NicheNet analysis showing the expression of ligands received by CD4 T cells (compared to Naive) in the sender cell type after each route of vaccination.

### Activated IM localise in distinct tissue niches within the lung

Since NicheNet had indicated that IM and CD4 T cells may be potential interaction partners, we decided to investigate if these cell types localized to distinct niches within the tissue using spatial transcriptomics. Lung slices were taken 8 weeks after IV, IT or SC BCG, as well as from naïve controls, and analyzed using the 10x Genomics Visium platform. Spatial clustering resulted in 21 clusters across the samples (Fig. 5 A, Supplementary Fig. 5 A). Of these, clusters 4, 18 and 19 were found predominately in IT and IV slices but not in SC BCG or naïve samples (Fig. 5 B, C, Supplementary Fig. 5 B). To determine the cellular composition of these vaccination induced clusters, we performed cell type deconvolution using RCTD based on our parallel single cell data. This analysis returns weights that estimate the cell type proportions in each spot. Cluster 4 had a bigger proportion of structural cells such as endothelial cells and fibroblasts while cluster 18 and 19 were enriched with immune cells. In particular cluster 18 had a higher proportion of activated IM, B cells, T lymphocytes and other immune cells (Fig. 5 D).

**Figure 5:**
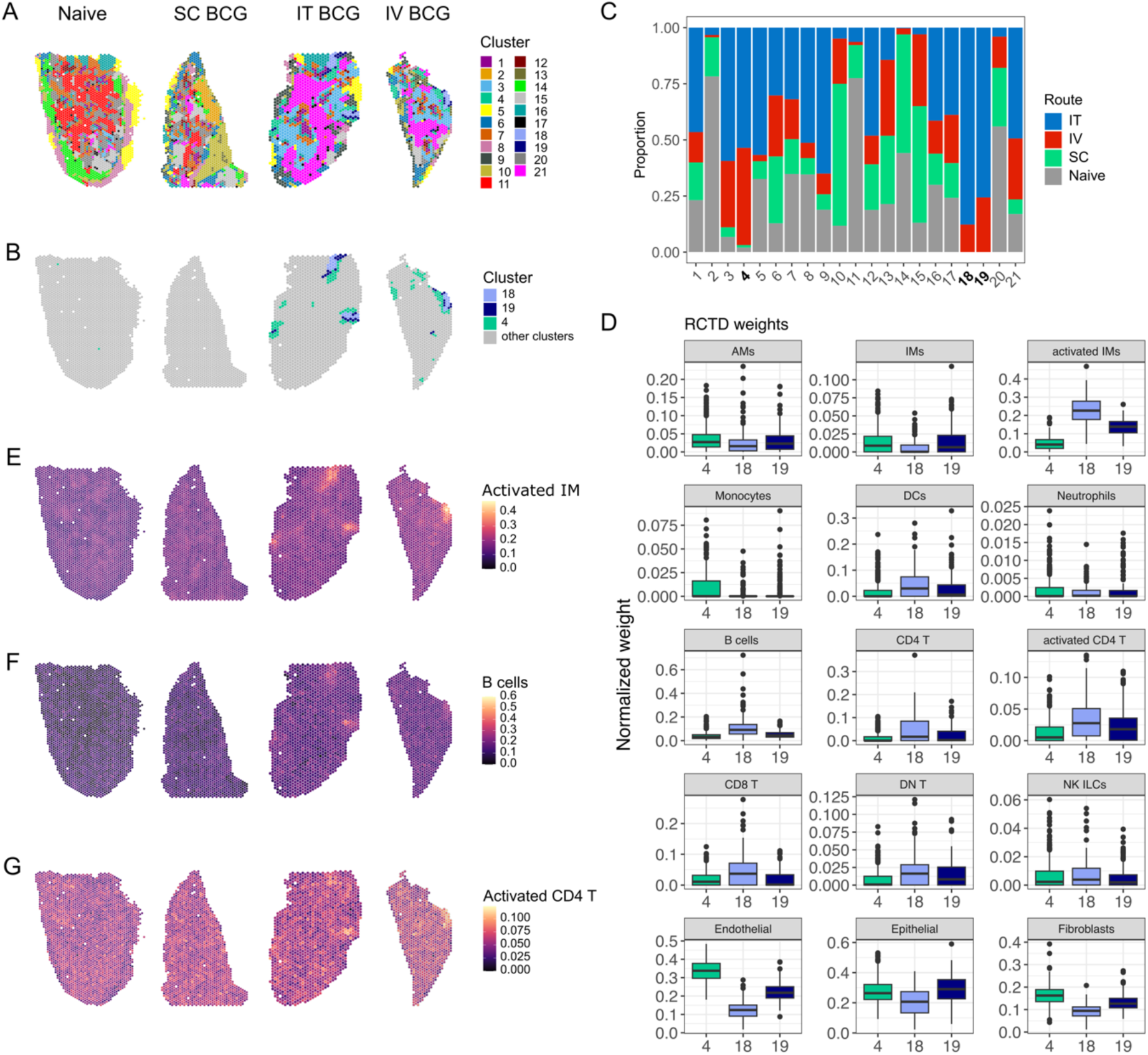
Activated IM localise in distinct tissue niches within the lung. **(A):** Spatial images depicting the clusters identified in naïve mice and animals that had been vaccinated 8 weeks previously. **(B):** Spatial images depicting only clusters 4, 18 and 19 in IT and IV BCG vaccinated samples. **(C):** The proportion of each cluster following each route of vaccination as well as in naive conditions. **(D):** RCTD weight plots of the immune and non immune cells present in cluster 4, 18 and 19. **(E-G)**: Overlay of activated IM (E), B cell (F) and activated CD4 (G) expression signature scores onto spatial slices taken from the lung of mice vaccinated 8 weeks prior as well as from naive animals.

To assess whether the activated immune cell populations identified in our scRNA-seq analysis localized to IT/IV specific tissue regions we overlaid our activated IM gene expression signature score onto the spatial slices. This signature was highly expressed in the regions occupied by spatial clusters 18 and 19 in the slices taken from IT and IV vaccinated lungs (Fig. 5 E, Supplementary Fig. 5 C). Similarly, B cell and activated CD4 T cell signatures showed strongest expression in these same tissue regions, with the CD4 signature showing somewhat broader distribution across the section (Fig. 5 F, G, Supplementary Fig. 5 D, E). Notably, expression of *Cxcl16* and *Cxcr6* was also concentrated in these areas, consistent with our NicheNet predictions of IM-CD4 T cell interactions (Supplementary Fig. 5 F). The co-localization of activated IM, B cells and CD4 T cells suggest that these regions are likely inducible bronchus-associated lymphoid tissue (iBALT), preferentially formed after IT and IV, but not SC, BCG vaccination. Whether proximity to iBALT facilitates rewiring of lung macrophages following BCG vaccination is currently unclear and warrants further investigation.

### IT BCG vaccination confers superior protection during Mtb challenge

To assess if the observed changes in IM phenotype and tissue localization following IT BCG translated to superior protection we performed rechallenge experiments. WT mice were vaccinated with BCG Pasteur via SC, IV, or IT routes. Sixty days later, vaccinated and unvaccinated mice were challenged with fluorescent Mtb, and lung bacterial burden was measured 14 days post-infection (Fig. 6 A). As compared to unvaccinated controls, mice that had received prior IT vaccination exhibited the greatest reduction in Mtb burden (Fig. 6 B). Flow cytometric analysis confirmed Mtb infection of neutrophils, DCs, AM, and IM, and revealed decreased Mtb signal in these populations in all vaccinated groups (Fig. 6 C and Supplementary Fig. 6 A-D).

**Figure 6:**
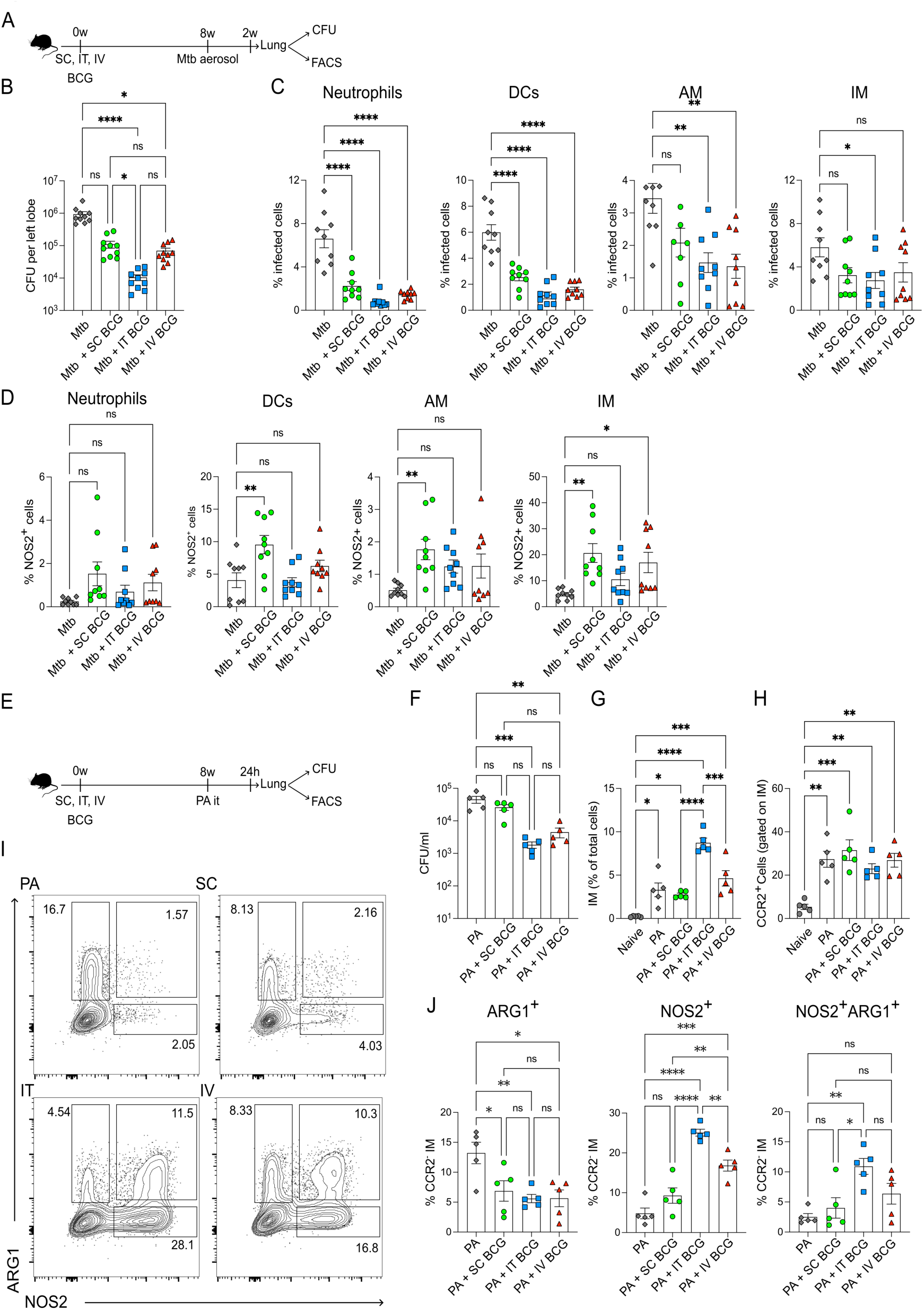
IT BCG vaccination confers superior protection during Mtb challenge. (**A):** Schematic depicting the experimental set up of the Mtb rechallenge. **(B)**: Quantification of Mtb bacterial burden 2 weeks after aerosol infection in unvaccinated mice or in animals that had received SC, IT or IV BCG vaccination 8 weeks previously. **(C):** Frequency of infected neutrophils, dendritic cells, AM and IM. **(D):** Bar plots depicting frequency of NOS2 positive neutrophils, DCs, AM and IM. **(E)**: Schematic depicting the experimental set up of the PA rechallenge. **(F)**: Quantification of PA burden 24 hours after infection in unvaccinated mice or in animals that had received SC, IT or IV BCG vaccination 8 weeks previously. **(G)**: Frequency of IM following PA infection as in (F). **(H)**: Quantification of CCR2 positive IM following PA infection. **(I)**: Representative FACs plots depicting Arg1 positive, NOS2 positive or double positive CCR2 negative IM. **(J)**: Quantification of Arg1 positive, NOS2 positive and double positive IM after PA infection. N: Each datapoint indicates one individual mouse. For B: One-Way ANOVA with non-parametric Kruskal-Wallis Test was performed. For C-J: One-Way ANOVA with Tukey’s multiple comparison test was used to determine statistical significance.

Next, we analyzed the expression of the anti-microbial effector molecule NOS2 across myeloid populations. Among the analyzed cell types, DCs - but not neutrophils - expressed NOS2, and the frequency of NOS2+ DCs increased following SC, but not IT or IV BCG (Fig. 6 D Supplementary Fig. 6 E, F). Very few AM exhibited NOS2 expression in Mtb infected mice and was only slightly enhanced by prior BCG vaccination (Fig. 6 D). Instead, robust NOS2 expression was observed in IM following Mtb infection (Fig. 6 D and supplementary Fig. 6 H). Notably, IM from SC vaccinated mice also expressed significant NOS2 levels after Mtb challenge, despite SC BCG having minimal impact on IM phenotype. Given the higher bacterial burden in SC vaccinated mice, this might reflect infection-driven rather than vaccine-induced NOS2.

To complement the above data and to more directly assess whether IT BCG enhanced IM effector function in the absence of antigen-specific adaptive immunity, we next established an acute infection model using the opportunistic pathogen *Pseudomonas aeruginosa* (PA). Cohorts of WT mice were vaccinated for 8 weeks as described above and then challenged with PA. 24 hours after infection bacterial burden in the lung was assessed via agar plating (Fig. 6 E). Mice that had received prior IT and IV, but not SC, BCG had significantly reduced bacterial load as compared to unvaccinated mice (Fig. 6F). Using flow cytometry, we observed that the frequency of IM increased after infection and this was most pronounced in mice vaccinated with IT BCG (Fig. 6 G). Additionally, CCR2 expression was increased on IM across all vaccination routes, likely indicative of monocyte influx following PA infection (Fig. 6H). Within CCR2- IM we observed only minimal NOS2 production from unvaccinated mice infected with PA. Instead, these cells expressed Arginase 1 (Arg1), a marker typically associated with the resolution stages of inflammation (Fig. 6 I, J). PA-induced Arg1 expression may indicate that IM are being skewed away from bacterial control and toward limiting tissue inflammation. Strikingly, prior IT BCG vaccination significantly boosted NOS2 expression by IM while limiting Arg1 (Fig. 6 I, J). Interestingly, we also observed an accumulation of NOS2+Arg1+ double positive IM in IT BCG mice infected with PA (Fig. 6 I, J). NOS2 and Arg1 expression have traditionally been used as markers of opposing macrophage polarization states, particularly in *in vitro* studies. The presence of an IM population co-expressing both markers suggests macrophage heterogeneity or the concurrent activation of multiple signaling pathways, underscoring the complexity of the inflammatory tissue microenvironment. Collectively our data demonstrate that mucosal BCG administration is protective against both homologous and heterologous bacterial pathogens and point to IM as a central mediator of protection.

## Discussion

In this study we investigated how the route of BCG vaccination differentially programs pulmonary macrophages and influences subsequent protection. Using high dimensional flow cytometry, we observed that IT and IV BCG vaccination resulted in similar upregulation of surface and activation markers in AM and IM, while macrophages from SC BCG mice more closely resembled naïve cells. ScRNA-seq confirmed that SC BCG induced minimal changes in AM and IM. Notably, our transcriptome data revealed that mucosal IT BCG predominantly rewired lung IM while exerting a less pronounced effect on AM. IM from IT BCG vaccinated mice exhibited enhanced expression of genes downstream of type I/II interferon signaling, as well as metabolic genes involved in glycolysis and oxidative phosphorylation. Our antibiotic data suggest that this IM phenotype is maintained without viable BCG, suggesting potential epigenetic rewiring following vaccination. Future work will incorporate challenge experiments in antibiotic-treated, BCG vaccinated mice to investigate this further.

Because IM functional capacity may depend on developmental origin, an unresolved question is whether activated IM induced after IT BCG arise from recruited monocytes or from locally rewired resident IM. This could be addressed by flow cytometric analysis of bone marrow to determine whether mucosal BCG alters progenitor composition, combined with genetic fate mapping models to define IM ontogeny. Consistent with systemic progenitor reprogramming, a non-human primate (NHP) study found that mucosal BCG induces transcriptional and metabolic signatures of trained immunity in blood and bone marrow monocytes (25).

Beyond IM reprogramming, our data further reveal spatially organized IM-CD4 T cell crosstalk as a potential mechanism underlying mucosal vaccine efficiency. Our transcriptome data identified CD4 T cells to be a source of IFNγ, and cell-cell communication analysis suggested reciprocal signaling between IM and CD4 T cells following IT BCG. Mucosal vaccination led to the enrichment of lung-resident CD4 T cells with increased expression of activation markers and chemokine receptors. Spatial transcriptomics further indicated that activated IM localize in close proximity to CD4 T cells within iBALT regions, raising the possibility that IT BCG establishes immune niches that position CD4 T cells and IM for sustained, bidirectional signaling. A previous study demonstrated that a subset of IM is required for iBALT formation in the lung during a house dust mite model (8). However, whether proximity to iBALT is necessary to drive IM activation—particularly during secondary infections—remains unclear. Conceptually, the iBALT niche may facilitate IM–CD4 T-cell crosstalk or provide additional cytokine signals beyond IFNγ that influence IM behavior. Experiments that disrupt iBALT formation, e.g. by using LTBR-Fc (26), after BCG vaccination but prior to secondary infection may shed light on this issue.

These spatial findings raise the possibility that IM guide CD4 T cell localization to and/or within the lung via CXCL16/CXCR6, which in turn facilitates IM activation via T cell derived IFNγ. Supporting this idea, a recent study found CXCL16-positive macrophages situated in close proximity to CXCR6-expressing T cells at the iBALT periphery following influenza infection (27). Future work to assess cytokines in lung homogenates across multiple timepoints post vaccination, as well as the effects of CXCL16 blockade and lymphocyte depletion, will help clarify the mechanistic basis of IT BCG induced programming of IM and the requirement for CD4 T cells in this process (28).

In our Mtb challenge model, mice vaccinated with IT BCG demonstrated the strongest protection (i.e. had the lowest bacterial burden) compared to the other routes. This increased protective capacity was not solely attributable to BCG presence in the lung, since IV mice had similar BCG burdens at 4- and 8-weeks post vaccination. We propose that direct IT delivery to the lung facilitates faster uptake of BCG by IM combined with earlier CD4 T cell recruitment, both of which would enable IM epigenetic adaptation. A functional consequence of IM reprogramming is further implied by our *P. aeruginosa* acute infection model, where IM from IT BCG vaccinated mice expressed the highest levels of NOS2. This suggests that IT BCG licenses IM for rapid activation after challenge, thereby facilitating early bacterial clearance. However, while our data strongly implicate IM as key mediators of IT BCG induced protection, we cannot exclude a contributory role for AM, particularly during early pathogen encounter. Future work using recently developed IM-specific (8,9) and AM-specific (29) depletion models will definitively address the relative contributions of each population during bacterial challenge.

Our findings strengthen the rationale for mucosal BCG as a vaccination strategy and complement recent controlled human infection model studies evaluating the effects of route-specific BCG vaccination in humans (30,31). However, our observation that SC and IV vaccinated mice exhibited similar Mtb burdens appears to diverge from the findings of Darrah et al. (16) who reported superior protection following IV BCG compared to intradermal administration in NHPs. Several factors may contribute to this difference. First, the kinetics and magnitude of BCG induced immunity may differ between mice and NHPs. Second, we assessed Mtb burden at 2 weeks post challenge whereas Darrah et al. evaluated protection at 12 weeks; accordingly, it is possible that IT and IV vaccination would both demonstrate superior protection relative to SC vaccination at later time points in mice. Third, differences in BCG inoculum (1 × 10^6^ in our study versus 5 × 10^7^ in Darrah et al.) and/or the dose for Mtb challenge (100-200 CFU in our study versus 4-36 CFU in Darrah et al.) may influence the magnitude of protection.

In summary, our study demonstrates that the route of BCG vaccination critically shapes pulmonary macrophage reprogramming, with mucosal delivery uniquely licensing IM for enhanced activation potentially through spatially organized immune interactions. These findings establish a mechanistic framework for optimizing mucosal vaccination strategies against tuberculosis and other respiratory pathogens.

## Methods

### KEY RESOURCES TABLE

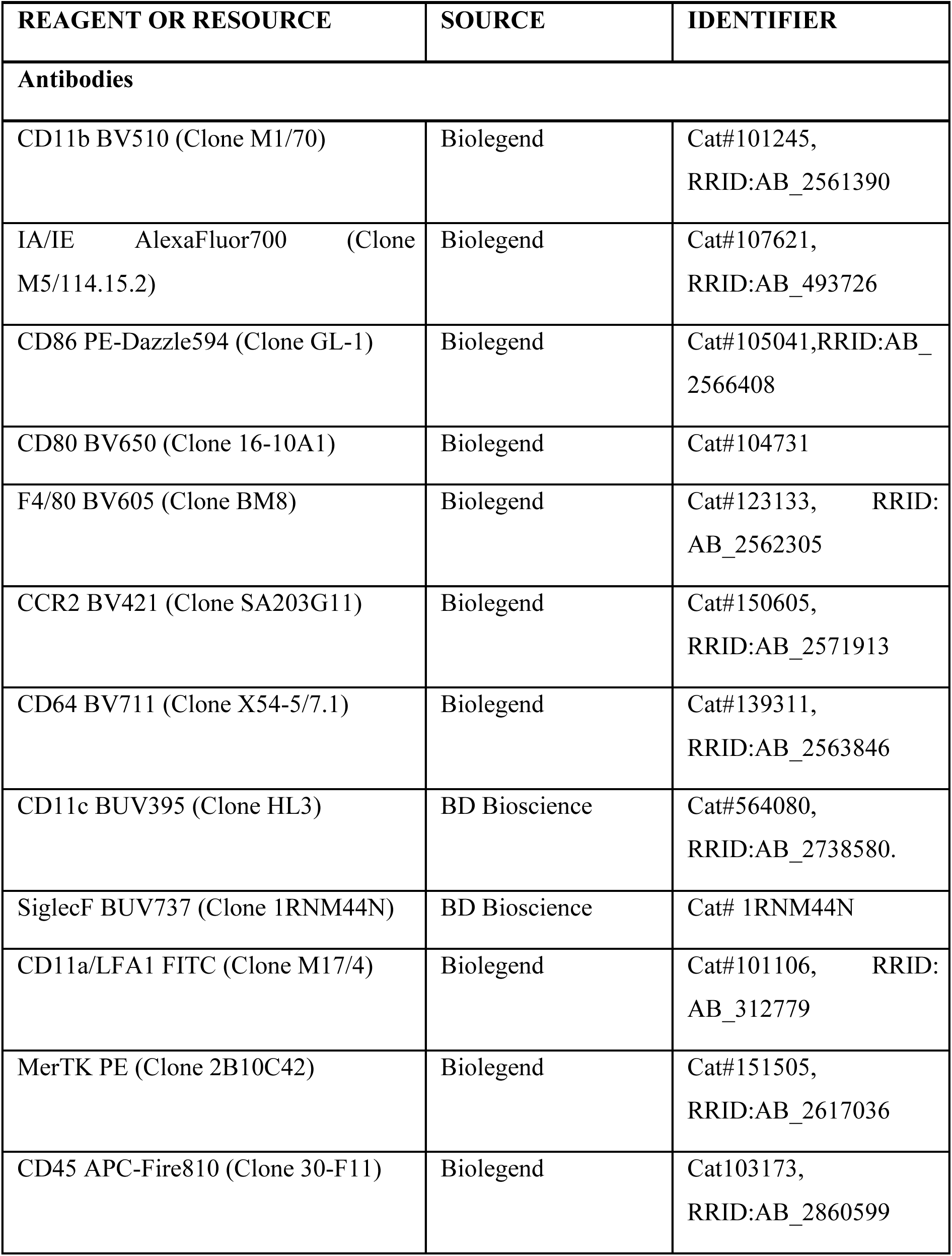
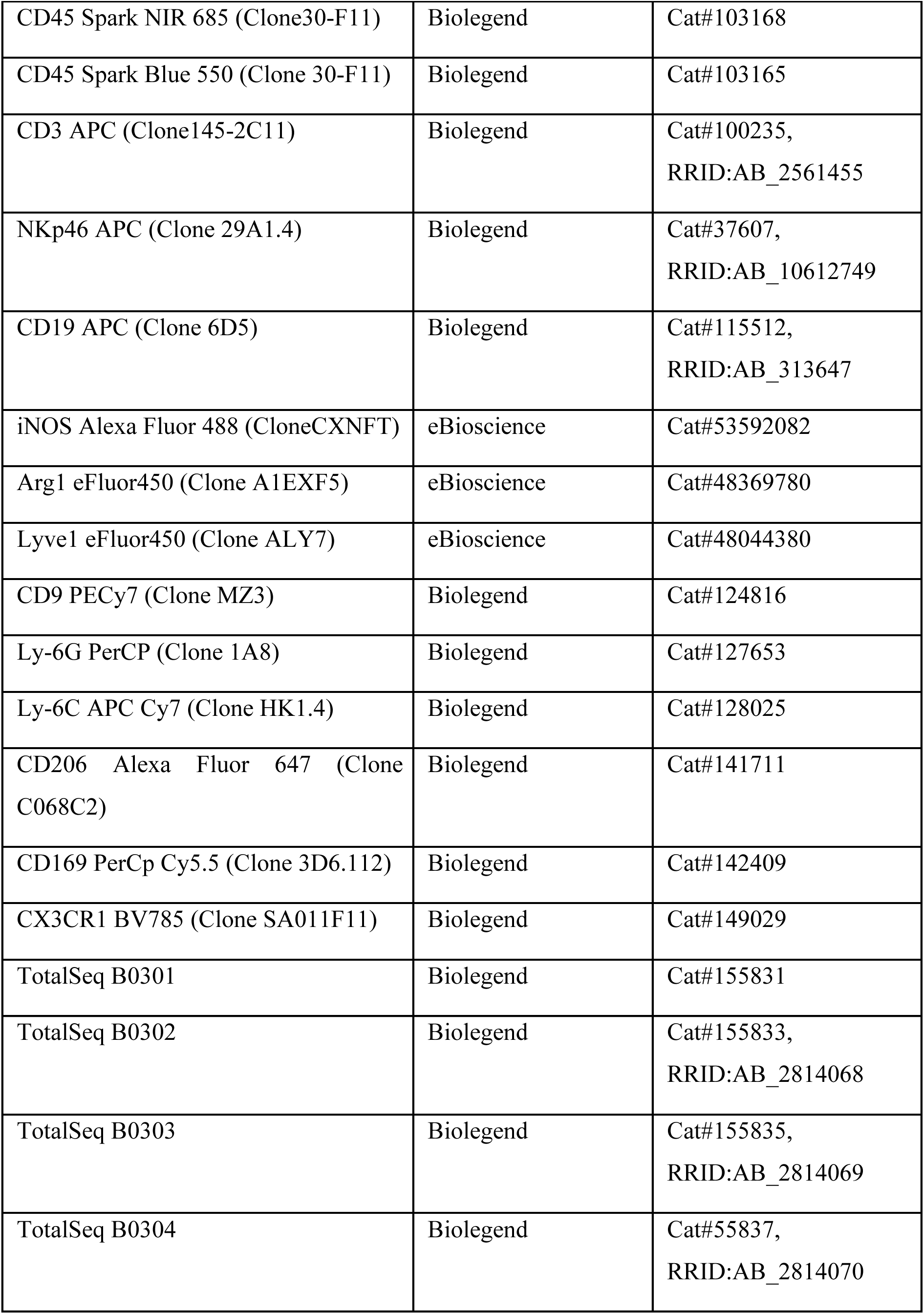
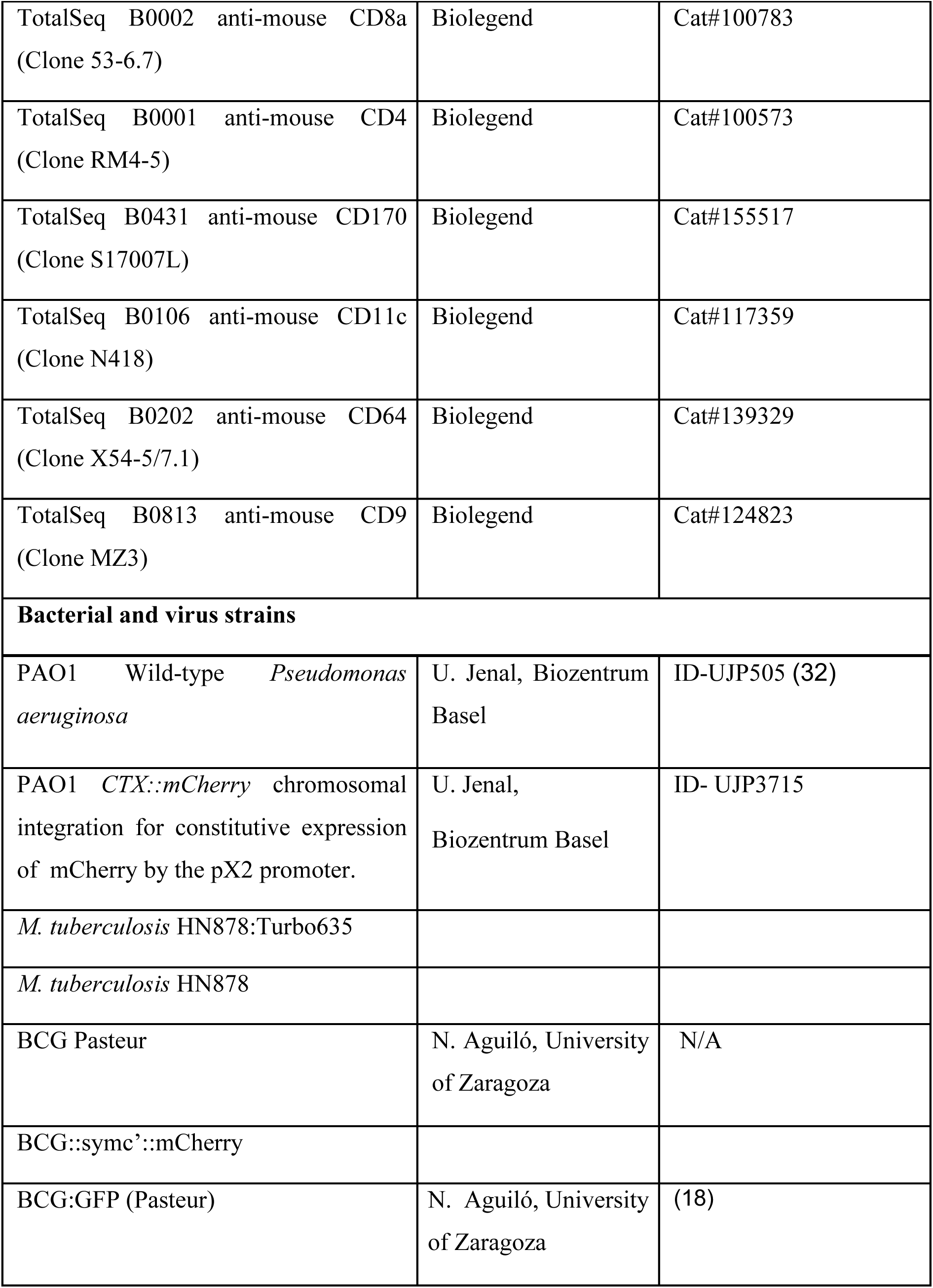
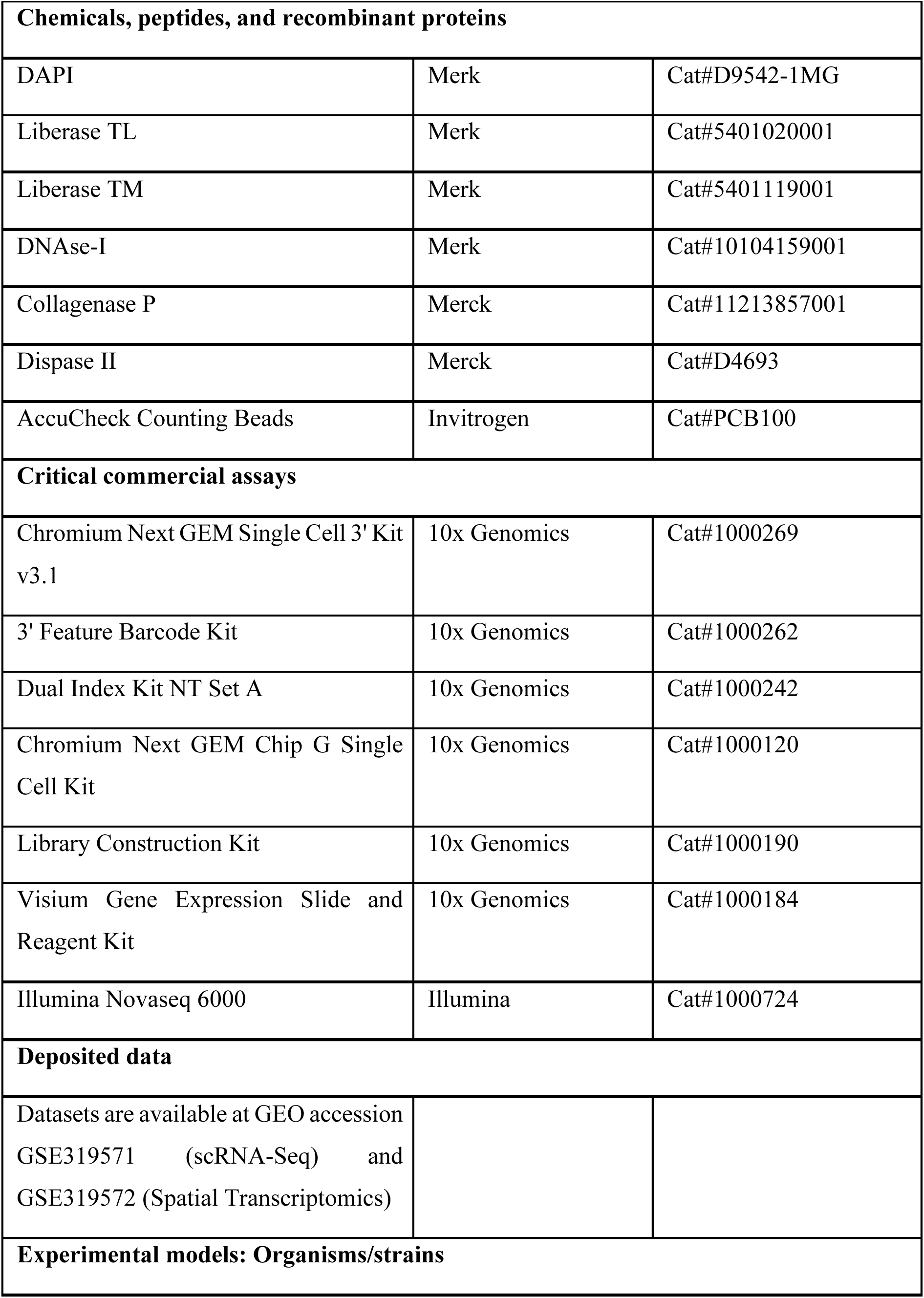
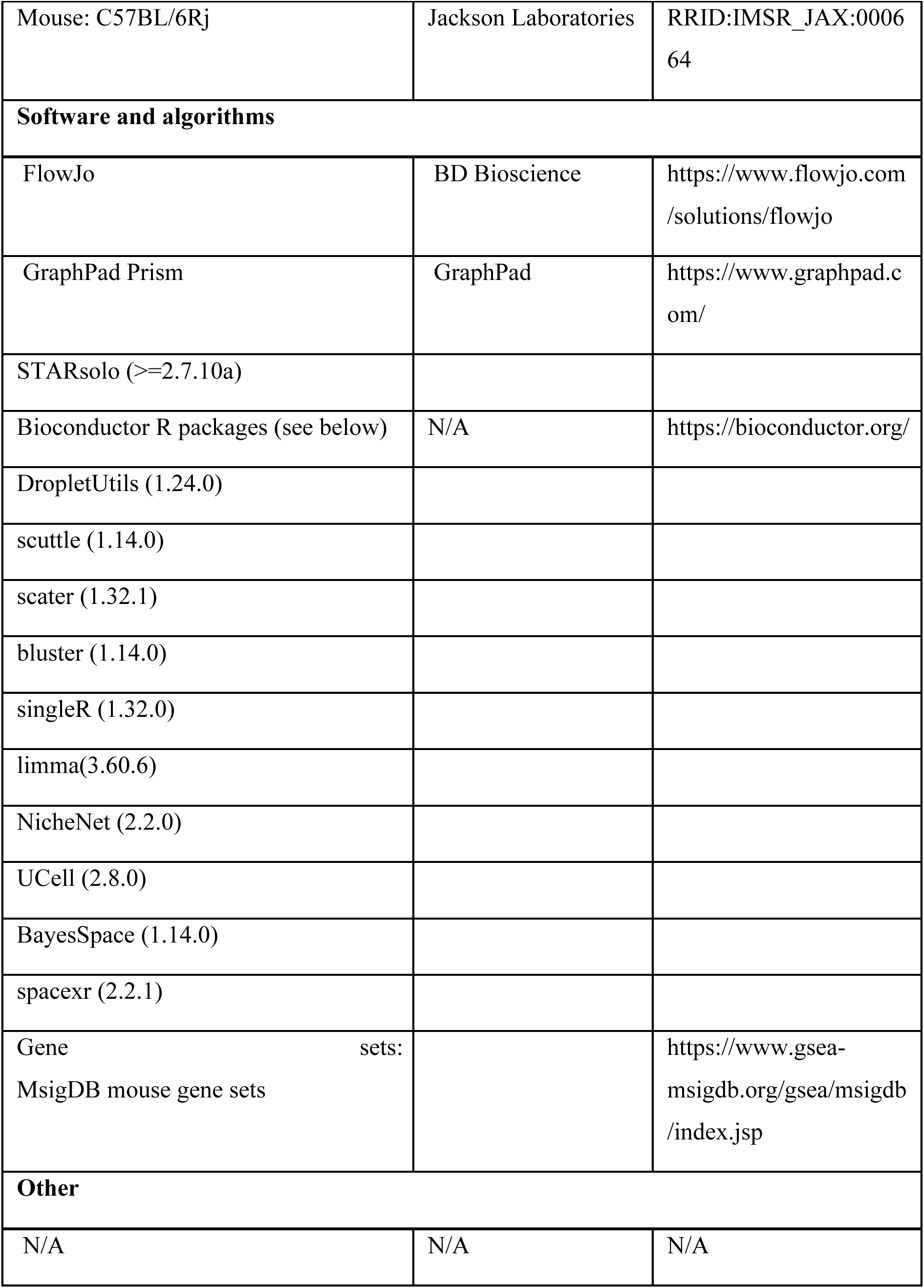

#### Animals

C57BL/6Rj (CD45.2) were purchased from Charles River. Mice in each experiment were 6-8 weeks old, same-sex littermates, maintained and bred in the specific pathogen–free animal facility at the University of Basel. All animal experiments were performed in accordance with local and Swiss federal guidelines.

#### Bacterial culture

*M. bovis* BCG::symc’::mCherry (hereafter BCG::mCherry) was generated in the laboratory by transformation with the pCHERRY3 plasmid (a kind gift from D. Russell, Cornell University). BCG::mCherry*, M. bovis* BCG Pasteur and BCG:GFP (kindly gifted by N. Aguiló University of Zaragoza) were grown in 7H9 broth (BD) 10% OADC (BD), 0.2% (v/v) glycerol and 0.05% (v/v) Tween 80 (Sigma) under constant shaking at 37° C. After 2 to 4 days the preculture was then diluted 1:100 in 7H9 media without Tween80. When bacteria reached an optical density (OD600) of 0.8 to 1, bacteria were collected and washed with PBS (Sigma, D8662). Subsequently bacteria was de-clumped by manual shaking with glass beads and the upper phase containing the single cell suspension was collected. De-clumped bacterial cultures were frozen in 25% glycerol at -80°C until further use. For colony-forming unit (CFU) determination bacterial suspension was plated on 7H11 agar plates.

HN878:turbo635 was generated in house by transformation with the plasmid pCHARGE3 kindly gifted by Tanya Parish (33) (Addgene plasmid # 24658; http://n2t.net/addgene:24658; RRID:Addgene_24658). HN878:turbo635 was grown in 7H9 broth (BD) 10% OADC (BD), 0.2% (v/v) Glycerol, 0.1 mM sodium propionate and 50 mg/ml Hygromycin B (Roth) . Bacteria were de-clumped by vortex shaking with glass beads and the upper phase containing the single cell suspension was harvested and frozen at -80°C until use.

*P. aeruginosa* (PAO1, PAO1 CTX::mCherry) was kindly provided by U. Jenal, University of Basel. *P. aeruginosa* was cultured in liquid LB and subsequently frozen in 70% glycerol. Prior to infection *P. aeruginosa* was plated from frozen glycerol stock overnight at 37°C on LB agar plates. Single colonies were picked and grown overnight at 37°C under constant shaking (170 rpm). Overnight cultures were then diluted and collected at log phase (OD600 = 0.5).

#### *Mtb* infection

Prior to aerolization, Mtb was declumped with a 27G metal tip. Mice were infected in an Aero3G Third Generation Aerosolizer (Biaera). Stocks were titrated to result in a day 1 bacterial load of approximately 100 CFU.

#### BCG vaccination / *P. aeruginosa* infection

For intratracheal administration mice were anesthetized with vaporized isoflurane and than infected intratracheally with 1x10^6^ CFU diluted in 50 μL of PBS (Gibco). For subcutaneous injection 1x10^6^ CFU was delivered in 200 μL, while intravenous BCG (1x10^6^ CFU) was administered in 100 μL PBS.

#### CFU plating

##### Mtb

For determination of CFU the left lobe was mechanically homogenized in 1 mL of PBS-Tween (0.05%) using soft tissue homogenizing CK28 prep tubes (Bertin technologies) in a Precellys Evolution Touch homogenizer (Bertin technologies). Homogenates were then serially diluted and plated on 7H11 agar plates containing 10% OADC, 0.2% (v/v) Glycerol, 50 mg/ml Hygromycin B and 1 vial of MGIT PANTA™ Antibiotic mixture (BD). CFU were counted after a 3 week incubation at 37°C.

##### P. aeruginosa

For determination of CFU lung lobes were digested as described above. 50 uL of digest were removed prior to filtration and a serial dilution was plated on tetracycline selective LB agar plates. CFU were counted after overnight incubation at 37°C.

#### Tissue preparation

In some experiments, prior to euthanasia mice were anesthetized with vaporized isoflurane and a CD45 (Spark Nir 685) antibody was administered intratracheally. After 20 minutes mice were injected with CD45 (APC-Fire 810) intravenously and sacrificed after 5 minutes.

For isolation of macrophages or lymphocytes, lungs were harvested, minced with scissors and digested with a digestion mix containing liberase TM or liberase TL (500 μg/ml, Roche) for 30 minutes at 37° C under constant shaking. Single cell suspensions were generated by repeated pipetting and filtering through 70 μm cell strainers (Milteny Biotec) in HBSS buffer (Hanks’ Balanced Salt solution, Sigma, H6648-500mL, 0.2% BSA and 0.5 mM EDTA). Erythrocytes were lysed by addition of erythrocyte lysis buffer and the remaining lung cells were washed with HBSS buffer.

#### Single cell sequencing

For isolation of lymphocytes lungs were collected and diced into gentleMACS C Tubes (Miltenyi Biotec) and washed down with 3 ml of media (RPMI, 10 mM aminoguanidine hydrochloride (Sigma-Aldrich), 10 mM Hepes, penicillin-streptomycin-glutamine (100×, Gibco)), prewarmed in a 37° C water bath. Digestion mix containing liberase TL (for lymphoid cells) (33.3 μg/ml, Merk) and deoxyribonuclease I (58 μg/ml; Applichem) was added. Lungs were dissociated on a gentleMACS Dissociator (Miltenyi Biotec) and mashed through 70 μm MACS SmartStrainers (Miltenyi Biotec) as previously described (34). The isolation of stromal cells and myeloid cells lungs has previously been described (35). In short: lobes were minced with scissors and digested with 1mL of RPMI-1640 containing 0.8 mg/ml DispaseII and 0.2 mg/ml Collagenase P (both from Roche), and 0.1 mg/ml DNase I (Invitrogen) at 37° C on a heating block and gently inverted and mixed using a 1mL pipette every 5 min. After 15 min large fragments were spun down (10s) and the supernatant was removed and filtered through a 70 μm MACS SmartStrainer into a tube containing 5mL of Lung Media ((RPMI, 10 mM aminoguanidine hydrochloride (Sigma-Aldrich), 10 mM HEPES, penicillin-streptomycin-glutamine (100x, Gibco), 2% FCS, 0.5mM EDTA)). Another 1mL of digestion mix was added to the remaining lung fragments and the process was repeated two more times. Finally, erythrocytes were lysed by addition of ELB (155mM NH4Cl, 170mM tris-HCL), remaining cells were washed with FACS buffer (PBS, 2% FCS, and 0.1% sodium azide) and further processed for sorting and subsequent single cell sequencing.

#### Flow cytometry staining

For all fluorochrome-conjugated antibody dilutions, FACS buffer containing Fc block (Clone 2.4G2 InVivoMAb anti-mouse CD16/CD32; BioXCell, #BE0307) was used. Live-dead stain (Zombie Fixable Viability™ Sampler Kit; BioLegend) was added to the antibody mix and stained at RT 10 min. FoxP3 Transcription Factor Staining Buffer Set (eBioscience) was used for intracellular staining. Flow cytometric analysis was performed on Cytek Aurora. Data was analyzed using FlowJo X software (BD) and SpectroFlo (Cytek). Absolute cell numbers were determined using AccuCheck Counting Beads (Invitrogen) based on the manufacturer instructions.

#### Single cell RNA sequencing

Lungs of naïve or BCG infected mice were removed and processed as described above for isolation of lymphocytes or stromal/myeloid cells. Individual mice were hash tagged using TotalSeq-B anti-mouse hashtag antibodies (B0301-B0304, BioLegend) and TotalSeq-B CITE-seq antibodies (lymphocytes: CD8a (B0002), CD4 (B0001); myeloid cells: SiglecF (B0431), CD11c (B0106), CD64 (B0202), CD9 (B0813)). Live, CD45+ and CD45- cells were sorted from lungs at 80 d.p.i. and submitted for library preparation. A total of 10,000 target cells per sample were loaded. Single-cell capture and cDNA library preparation was performed according to manufacturer’s instructions using 10x Genomics Chromium Next GEM Single Cell 3’ Kit v2 reagents using dual indexing. Sequencing was carried out on an Illumina Novaseq 6000 at the Genomics Facility Basel, ETH Zurich.

Raw reads were mapped to the *Mus musculus* reference mm10_ensdb102 using STARsolo (36), and empty droplets were removed with DropletUtils (37,38). The four replicate samples were demultiplexed using hashtag oligos (HTO). Cells were filtered for library size, library complexity, and mitochondrial content using adaptive thresholding implemented in the *perCellQCFilters* function from the R package scuttle with default arguments (39). PCA and UMAP on the PCA-reduced dimensions, were performed using the R package scater (39). Cells were clustered either hierarchically using Ward’s method or with the graph-based Leiden method using the bluster R package. Cell annotation was performed with SingleR (40), and pseudo-bulk differential expression analysis was conducted using the *pseudoBulkDGE* function from the scran R package (1.32.0), following the Bioconductor OSCA book (https://bioconductor.org/books/release/OSCA/). Lists of differentially expressed genes per cell type and BCG route comparison are available in Supplementary Table 1. Gene set enrichment analysis was performed using camera (41) from the limma R package (42) with MSigDB mouse gene sets. Cell-cell interactions were interrogated using NicheNet (43) following the step-by-step vignette starting from a Seurat object. When possible, lists of differentially expressed genes derived from the pseudo-bulk differential expression analysis were provided to NicheNet for comparison across cell types. Gene signature scores were calculated using UCell (44).

#### Visium Spatial gene expression

Lungs were inflated with a 1:1 mixture of Tissue-Tek® O.C.T. Compound (Sakura) and PBS and embedded in O.C.T. Tissues were sectioned at 10 μm on a cryostat. Further processing and placement of sections was performed according to the protocol CG000240 (10X Genomics). Following sample placement, sections were stained with DAPI according to the protocol CG000312 (10X Genomics) and imaged using a Leica DMI 4000/6000 Widefield Microscope. Sections were then permeabilized and prepared for library preparation according to the protocol CG000239 (10X Genomics). Sequencing was carried out on an Illumina Novaseq 6000 at the Genomics Facility Basel, ETH Zurich.

The fiducial frame of the Visium slides was aligned using Loupe Browser, and the resulting JSON file was provided to Space Ranger to obtain count matrices. Spots outside the tissue, with library size < 2000, or with mitochondrial content > 10% were filtered out. Spatial clustering was performed using BayesSpace after joining spatial coordinates across samples and integrating transcriptomic data, enabling spatial clusters to be shared between samples (45). The single-cell dataset was used as a reference for deconvolution with RCTD from the spacexr R package (46). Signature scoring and plotting were performed as for single-cell data using Bioconductor 3.19.

#### Quantification and statistical analysis

Prism software (GraphPad 10. 4.1) was used for all statistical analyses. Unless otherwise stated One-Way ANOVA with Tukey’s multiple comparison test was used to determine statistical significance. Statistical significances are indicated by \**p*<0.05, \*\**p*<0.005, \*\*\**p*<0.0005, \*\*\*\**p*<0.0001, ns, not significant (*p* > 0.05). In summary graphs, points indicate individual samples. All error bars are mean with SEM.

## Acknowledgements.

We are indebted to Gaël Auray and all members of the University of Basel flow cytometry core facilities for cell sorting and troubleshooting, furthermore we thank all staff at the BSL2 and BSL3 animal facilities of the University of Basel. In addition we are grateful to the University of Basel Genomics Core. Bioinformatic calculations were performed at sciCORE (http://scicore.unibas.ch/) scientific computing center at University of Basel. We thank all members of the King laboratory for productive and helpful discussion. This work was supported by the Swiss National Science Foundation, grant 212344 awarded to CGK, the State Secretariat for Education, Research and Innovation (SERI), grant REF-1131.52304 awarded to CGK and the University of Basel Research Fund for Junior Researchers awarded to AJF. The authors have no conflicting financial interests.

---

**Supplementary Figure 1:**
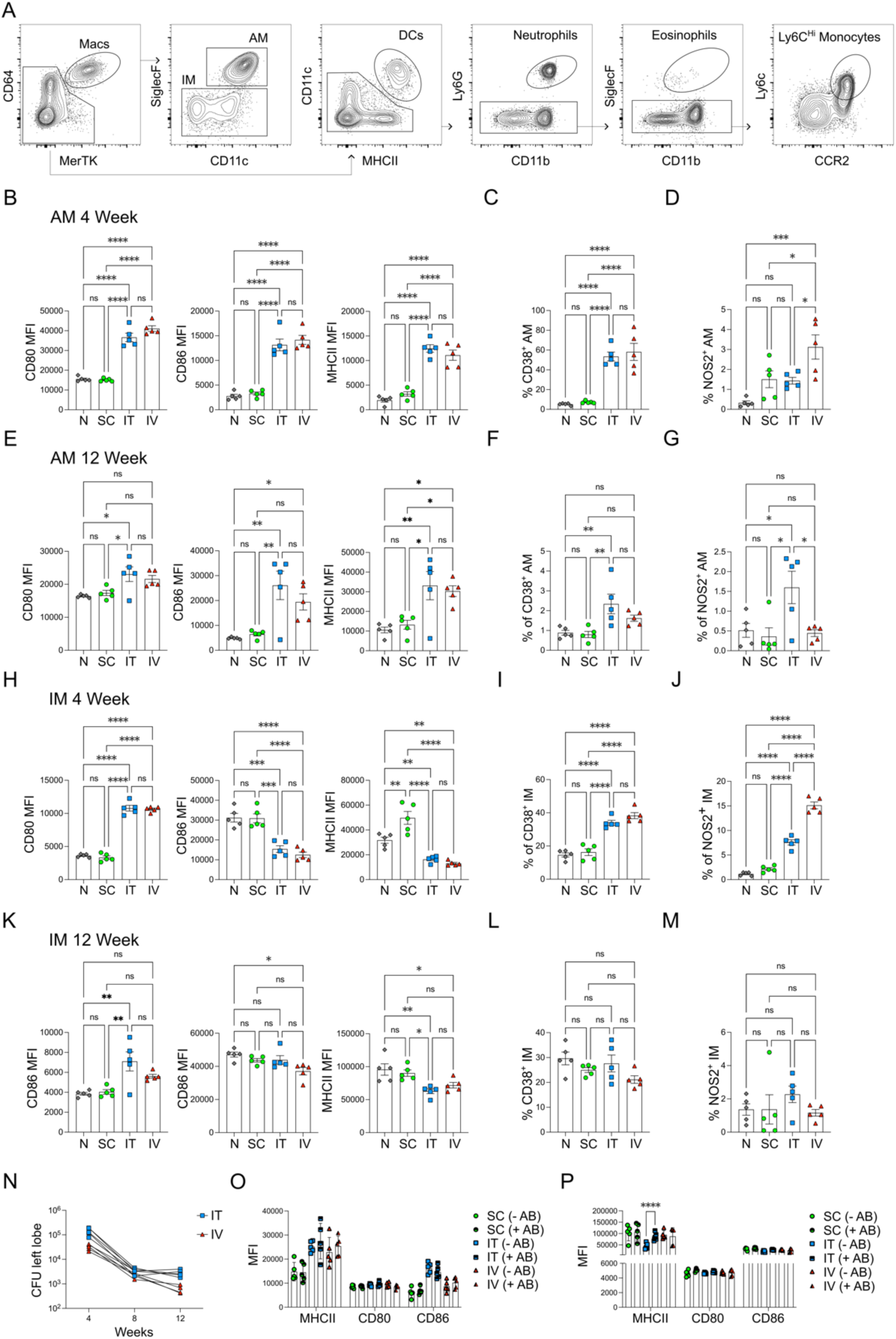
IT BCG supports enhanced IM activation. **(A)**: Gating strategy used to identify lung innate immune cells including AM, IM, DCs, neutrophils, eosinophils and Ly6C Hi monocytes. Gated on live, CD45+, CD3-, CD19- & Nkp46-cells. **(B)**: MFI of CD80, CD86 and MHCII expression by AM 4 weeks after SC, IT or IV BCG vaccination. **(C, D)**: Frequency of CD38 or NOS2 positive AM from mice vaccinated as in (B). **(E-G)**: MFI of CD80, CD86 and MHCII expression by AM 12 weeks after vaccination (E) and the frequency of CD38 or NOS2 positive cells (F, G). **(H)**: MFI of CD80, CD86 and MHCII expression by IM 4 weeks after SC, IT or IV BCG vaccination. **(I, J):** Frequency of CD38 or NOS2 positive IM from mice vaccinated as in (H). **(K-M)**: MFI of CD80, CD86 and MHCII expression by IM 12 weeks after vaccination (K) and the frequency of CD38 or NOS2 positive cells (L, M). **(N)**: BCG bacterial burden in the lungs of mice vaccinated with IT or IV BCG for 4, 8 and 12 weeks. **(O, P)**: MFI of MHCII, CD80 or CD86 expression by AM (O), or by IM (P), in mice that had been vaccinated with BCG and subsequently treated with antibiotics. N: Each datapoint indicates one individual mouse. One-Way ANOVA with Tukey’s multiple comparison test was used to determine statistical significance.

**Supplementary Figure 2:**
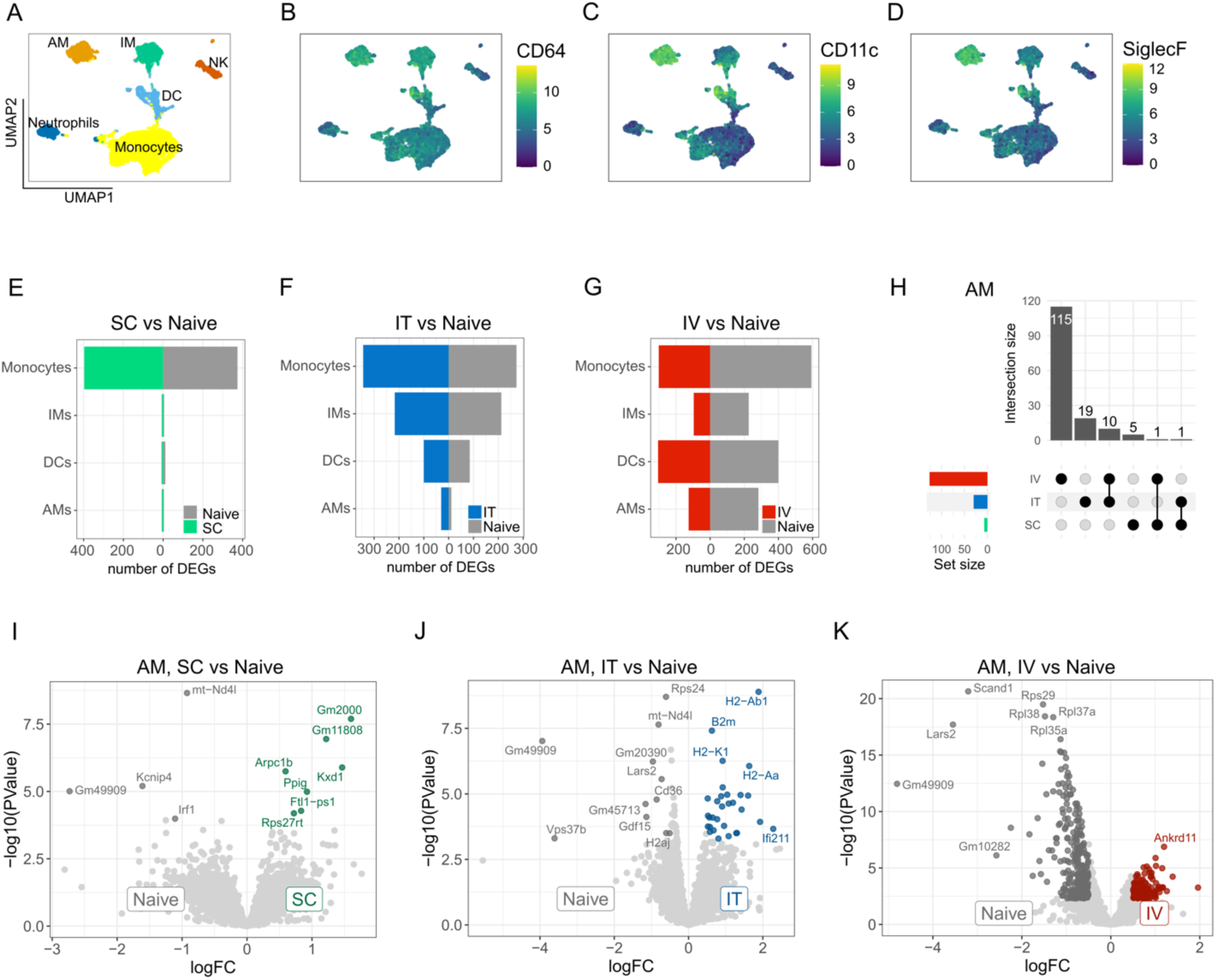
IT BCG transcriptionally rewires IM. **(A):** UMAP depicting clusters of AM, IM, DC, neutrophils and DCs identified in single cell RNA Seq dataset. **(B-D):** UMAP depicting protein expression of CD64 (B), CD11c (C) and SiglecF (B). **(E-G)**: Barplots depicting the number of differentially expressed genes in monocytes, DCs, IM or AM 8 weeks after SC, IT or IV BCG vaccination. **(H)**: Upset plot depicting the number of genes upregulated by AM after each route of BCG vaccination as compared to naïve cells. **(I-K)**: Volcano plots depicting genes expressed by AM following SC (I), IT (J) or IV (K) vaccination. Coloured dots represent significantly differentially expressed genes with FDR < 0.05 and |logFC| > 0.5 (dark grey for Naive, green for BCG SC, blue for BCG IT, red for BCG IV). Light grey dots represent non-significant genes.

**Supplementary Figure 3:**
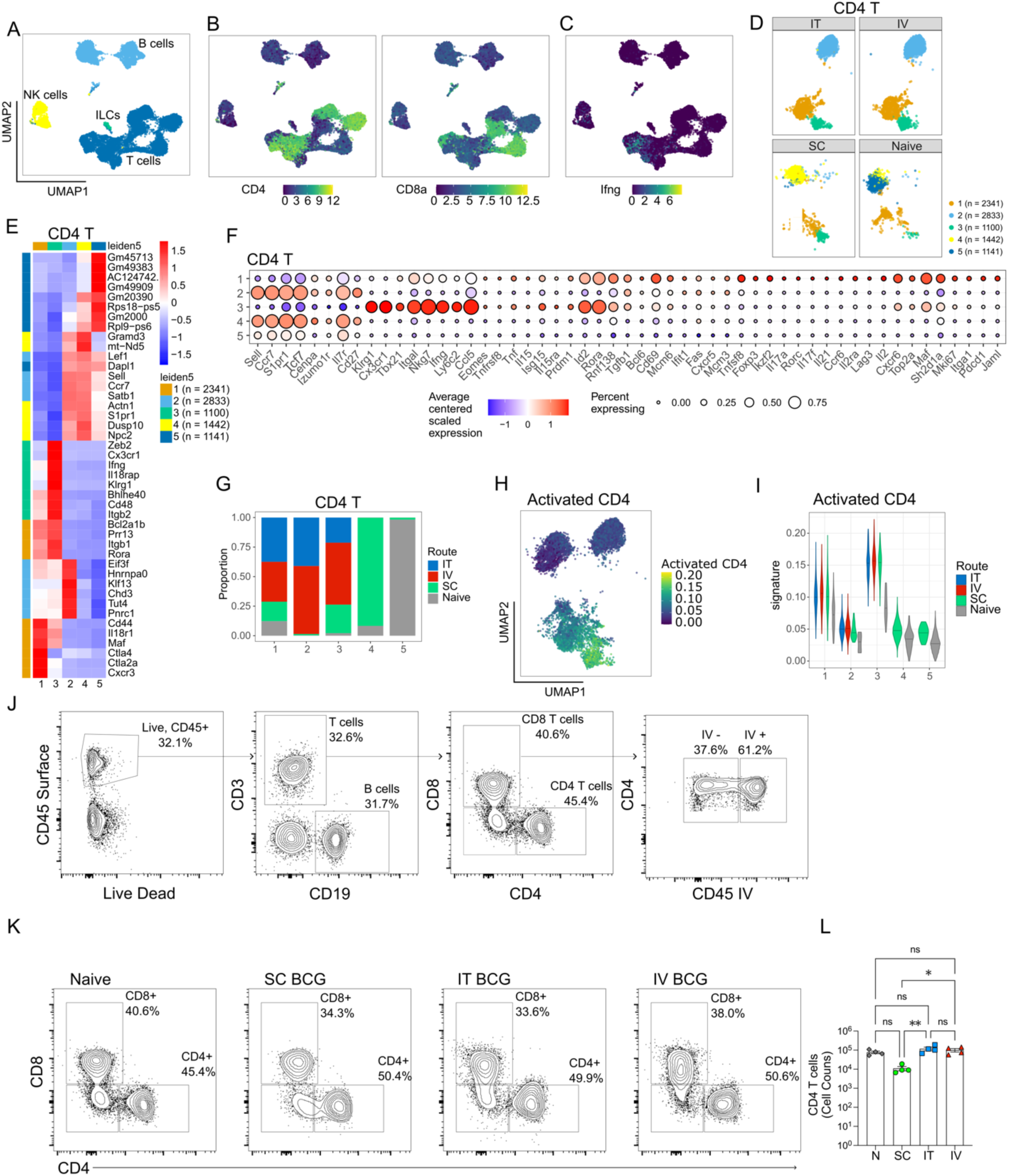
CD4 T cells likely produce IFNγ after all routes of BCG vaccination. **(A, B):** UMAP depicting lymphocyte populations identified in single cell RNA Seq (A) and Protein expression of CD4 and CD8 (B). **(C):** Expression of IFNgamma by the lymphocyte populations identified in (A). **(D)**: Clusters of CD4 T cells identified 8 weeks after SC, IT and IV BCG vaccination. **(E, F)**: Heatmap with the top 10 genes expressed by each cluster of CD4 T cells (E) and the expression of marker genes (F) used to identify the CD4 subsets. **(G)**: Proportion of each cluster of CD4 T cells 8 weeks after SC, IT and IV BCG vaccination. **(H)**: Expression of the activated CD4 signature score on the clusters of CD4 T cells. **(I)**: Violin plot depicting the strength of the activated CD4 expression signature in each CD4 T cell cluster. **(J)**: Gating strategy used to identify CD4 T cells. **(K, L)**: Representative plots and cell counts of CD4 T cells in naive and BCG vaccinated mice. For L, N: Each datapoint indicates one individual mouse. One-Way ANOVA with Tukey’s multiple comparison test was used to determine statistical significance.

**Supplementary Figure 4:**
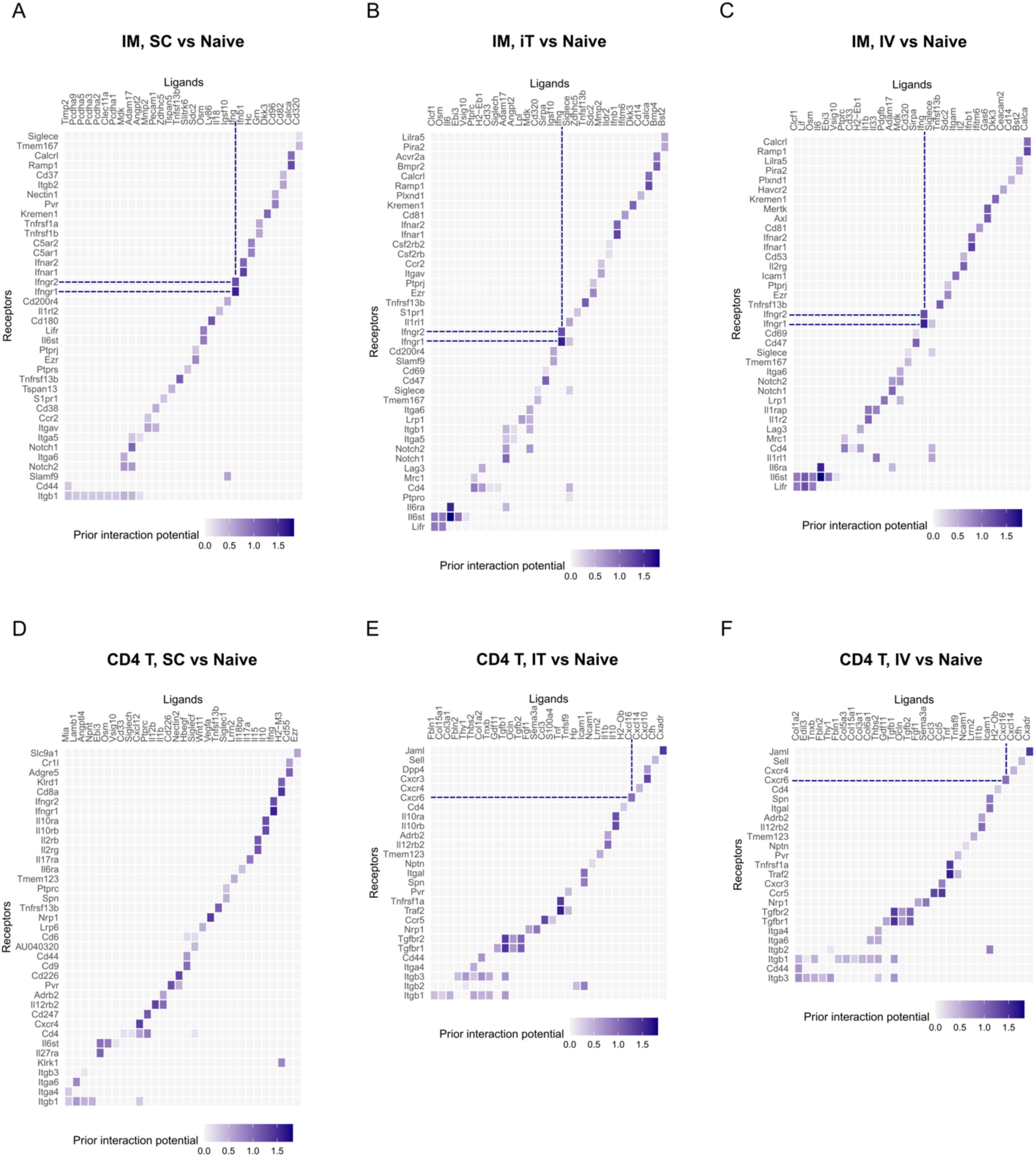
IM-CD4 T cell crosstalk following BCG Vaccination. **(A):** Heatmap representing NicheNet-predicted interaction potentials for ligand-receptor pairs, between the sender cell types and IM, that are contributing to downstream differentially expressed genes identified comparing BCG SC vs Naïve, BCG IT vs Naïve **(B)** and BCG IV vs Naïve **(C)**. The interaction between *Ifng* and *Ifngr1*/*Ifngr2* is highlighted. **(D):** Heatmap representing NicheNet-predicted interaction potentials for ligand-receptor pairs, between the sender cell types and CD4 T cells, that are contributing to downstream differentially expressed genes identified comparing BCG SC vs Naïve, BCG IT vs Naïve **(E)** and BCG IV vs Naïve **(F)**. The interactions between Ifng and *Ifngr1*/*Ifngr2* and between Cxcl16 and Cxcr6 are highlighted.

**Supplementary Figure 5:**
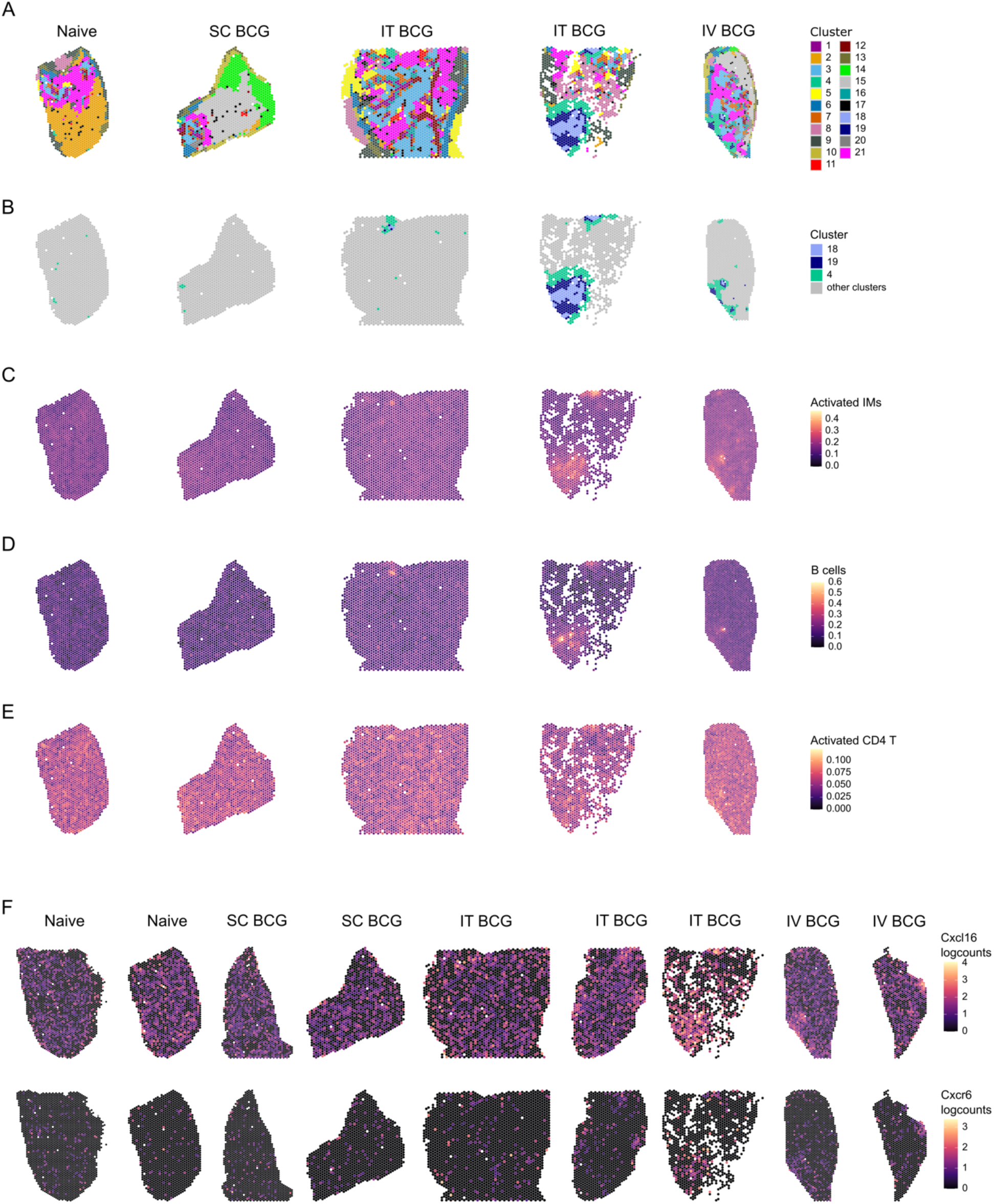
Activated IM localise in distinct tissue niches within the lung. **(A):** Spatial images of additional samples, depicting the clusters identified in naïve mice and animals that had been vaccinated 8 weeks previously. **(B)**: Spatial images of additional samples, depicting only clusters 4, 18 and 19 in IT and IV BCG vaccinated samples. **(C-E):** Overlay of activated IM (C), B cells (D) and activated CD4 T cells (F) expression signature scores onto additional spatial slices taken from the lung of mice vaccinated 8 weeks prior as well as from naive animals. **(F):** *Cxcl16* (top) and *CxCr6* (bottom) gene expression on all spatial slices taken from the lung of naive animals as well as from mice vaccinated 8 weeks prior.

**Supplementary Figure 6:**
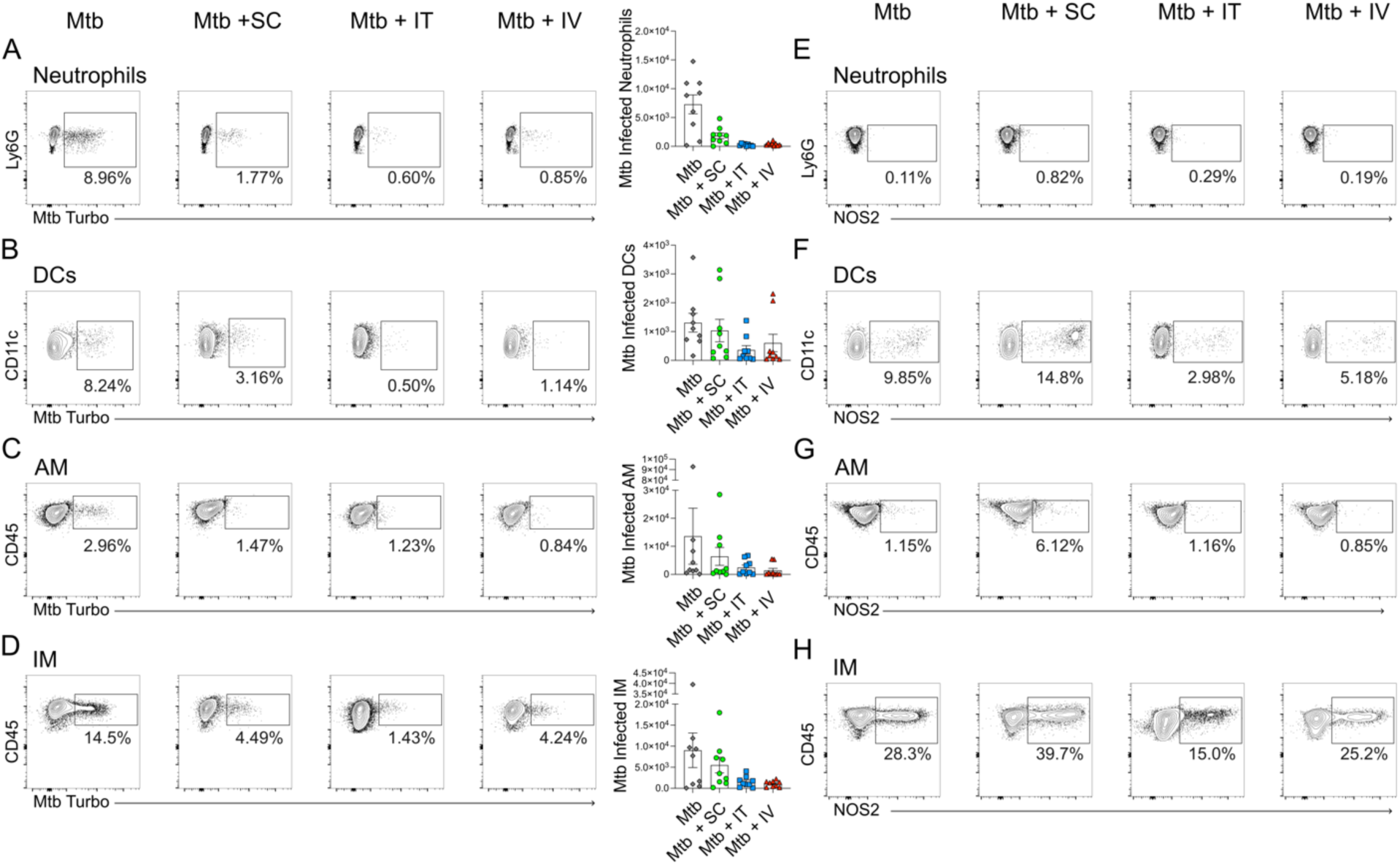
IT BCG vaccination confers superior protection during Mtb challenge. **(A-D)**: Representative FACS plots depicting Mtb-turbo signal in neutrophils (A), DCs (B), AM (C) and IM (D). Bar plots depict the number of infected cells for each condition. **(E-H)**: Representative FACs plots depicting NOS2 expression in neutrophils (E), DCs (F), AM (G) and IM (H).

